# Differential and synergistic effects of low birth weight and Western diet on skeletal muscle vasculature, mitochondrial lipid metabolism and insulin signaling in male guinea pigs

**DOI:** 10.1101/2021.10.13.464235

**Authors:** Kristyn Dunlop, Ousseynou Sarr, Nicole Stachura, Lin Zhao, Karen Nygard, Jennifer A. Thompson, Jennifer Hadway, Bryan S. Richardson, Yves Bureau, Nica Borradaile, Ting-Yim Lee, Timothy R.H. Regnault

**Author notes:** These authors contributed equally to this work (Kristyn Dunlop, Ousseynou Sarr). Kristyn Dunlop < >; Ousseynou Sarr < >; Nicole Stachura < >; Lin Zhao < >; Karen Nygard < >; Jennifer A. Thompson < >; Jennifer Hadway < >; Bryan S. Richardson < >; Yves Bureau < >; Nica Borradaile < >; Ting-Yim Lee < >; Timothy R.H. Regnault < >. Correspondence; Dental Science Building, Room 2021, Western University, 1151 Richmond Street, London, Ontario, N6A 5C1, Canada.

## Abstract

Low birth weight (LBW) offspring are at increased risk for developing insulin resistance, a key precursor in metabolic syndrome and type 2 diabetes mellitus. Altered skeletal muscle vasculature, extracellular matrix, amino acid and mitochondrial lipid metabolism, and insulin signaling are implicated in this pathogenesis. Using uteroplacental insufficiency (UPI) to induce intrauterine growth restriction (IUGR) and LBW in the guinea pig, we investigated the relationship between UPI-induced IUGR/LBW and later life skeletal muscle arteriole density, fibrosis, amino acid and mitochondrial lipid metabolism, markers of insulin signaling and glucose uptake, and how a postnatal high-fat, high-sugar “Western” diet (WD) modulates these changes. Muscle of 145-day-old male LBW glucose tolerant offspring displayed diminished vessel density and altered acylcarnitine levels. Disrupted muscle insulin signaling despite maintained whole-body glucose homeostasis also occurred in both LBW and WD-fed male ‘lean’ offspring. Additionally, postnatal WD unmasked LBW-induced impairment of mitochondrial lipid metabolism as reflected by increased acylcarnitine accumulation. This study provides evidence that early markers of skeletal muscle metabolic dysfunction appear to be influenced by the *in utero* environment and interact with a high fat-sugar postnatal environment to exacerbate altered mitochondrial lipid metabolism promoting mitochondrial overload.

## 1. Introduction

Metabolic syndrome (MetS) is an important risk factor for cardiovascular disease and type 2 diabetes mellitus (T2DM) with increased morbidity and mortality [1]. Insulin resistance is the primary determinant of the components of the MetS including obesity, glucose intolerance, dyslipidemia and hypertension and is evident before overt disease develops [1,2]. Insulin resistance develops in key insulin target tissues including muscle and is the result of altered fatty acid transport and oxidation, intramyocellular lipid accumulation and reduced mitochondrial oxygen uptake in skeletal muscle [3–5]. Insulin resistance is also characterized by altered insulin signaling responses associated with impaired vasculature, increased acylcarnitine contents, altered amino acid availability, and diminished glucose uptake in skeletal muscle [6–12].

The increasing prevalence of T2DM/insulin resistance has been widely reported to occur in conjunction with the increasing consumption of the energy-dense high-fat/high sugar diet or Western diet (WD) [13,14]. Prior to whole-body insulin resistance being detected, multiple organs develop insulin resistance, with muscle developing insulin resistance prior to other organs such as liver and white adipose tissue, with a cascade effect then occurring that is finally displayed as wholebody insulin resistance and glucose intolerance [15,16]. Early phases of insulin resistance development in skeletal muscle include extracellular matrix remodeling with increased physical barriers for insulin and glucose transport and decreased vascular insulin delivery [17]. Moreover, chronic consumption of a high-fat diet generates fatty acid oxidation rates in muscle that outpace the tricarboxylic acid cycle (TCA) leading to incomplete β-oxidation with intramitochondrial accumulation of acyl-CoAs and respective acylcarnitines [6,18]. This mitochondrial overload with acylcarnitines accumulation may then promote alterations in the phosphorylation status of insulin signaling intermediates as key mediators of the pathogenesis of skeletal muscle insulin resistance [6,19]. In conjunction with altered β-oxidation pathways, skeletal muscle amino acid availability is also impacted, all contributing to the development of skeletal muscle insulin resistance [8,18].

Animal and human studies now clearly show that conditions associated with the development of MetS can also be attributed to an adverse *in utero* environment such as observed with uteroplacental insufficiency (UPI)-induced intrauterine growth restriction (IUGR) [19–23]. IUGR resulting in low birth weight (LBW) is now recognized as a central programming factor in lifelong impairments in muscle development and metabolism [24]. Indeed, fewer capillaries per muscle fiber are found in LBW piglets [25], and in fetal IUGR sheep, skeletal muscle amino acid concentrations are altered indicating impairments in amino acid metabolism [26]. Additionally, gastrocnemius muscle of lean adult, glucose intolerant and insulin resistant male LBW rat offspring displays inefficient insulin-induced insulin receptor (IR), insulin receptor substrate-1(IRS-1), and Akt substrate of 160 kDa (AS160) phosphorylation, and impaired glucose transporter type 4 translocation in gastrocnemius muscle [27]. Metabolomic studies in blood of IUGR/LBW human neonates show abnormalities in amino acids and increases in acylcarnitine content [28,29] while that in blood of adult mice born LBW show altered β-oxidation with increased blood acetylcarnitine (C2) and short-chain acylcarnitines (C3-C5) [30], indicating mitochondrial overload due to incomplete fatty oxidation in later life, changes which have been associated with later life adult metabolic disease risk [31,32]. Furthermore, skeletal muscle in postnatal UPI-induced IUGR/LBW rats shows down regulation of insulin receptor beta (IRβ) and reduced phosphorylation of phosphoinositide 3-kinase (PI3K)/Protein kinase (AKT) at serine 473 (phospho-Akt^Ser473^) while that in young-adult men born LBW shows reduced insulin-signaling proteins [20], indicating disruption in insulin signaling [33,34]. Together, these studies highlight that skeletal muscle structure, metabolism and insulin signaling pathways are susceptible to the *in utero* environment and may underly persistent changes in the offspring associated with early adulthood development of pre-clinical markers of MetS.

Recently, evidence has suggested that *in utero* influences and secondary insults in postnatal life, such as excessive energy supply (e.g. WD) may have a synergistic effect, further exacerbating an already dysfunctional system and promoting an earlier more extreme onset of MetS [35]. In addition, there has recently been increased recognition of insulin resistance and T2DM among the pediatric population, attributed to a combination of genetic predisposition and environmental factors, namely sedentary lifestyle and energy-dense foods [36]. Therefore, the contribution of the *in utero* environment to alterations in muscle structure and metabolic function should not be ignored when assessing children born LBW and risk of developing metabolic dysfunctions such as insulin resistance and T2DM, at an earlier age. The Western diet is widely available in society, and given UPI is the most common *in utero* insult leading to IUGR/LBW [37,38], and high prevalence of IUGR/LBW infants [39] with increased metabolic disease risk later in life maybe greater than currently thought [40], hence the importance of understanding the specific developmental pathways to MetS. Here, we sought to identify alterations in skeletal muscle extracellular matrix, vasculature, amino acid and mitochondrial lipid metabolism, glucose uptake and insulin signaling following UPI-induced IUGR/LBW independently, and in conjunction with early postnatal exposure to a Western diet, in young adulthood.

## 2. Materials and Methods

### 2.1 Ethics statement

All investigators understood and followed the ethical principles as outlined by Grundy [41] and study design was informed by the ARRIVE guidelines [42].

### 2.2. Animal handling

Time-mated pregnant Dunkin-Hartley guinea pigs (Charles River Laboratories, Wilmington, MA, USA) where housed in a constant temperature (20°C) and humidity (30%) controlled environment with 12h light/dark cycle. All animals had *ad libitum* access to standard guinea pig chow (LabDiet diet 5025) and tap water. All pregnant guinea pigs underwent partial ablation of branches of uterine artery to induce uteroplacental insufficiency and intrauterine growth restriction (IUGR) [43]. At mid-gestation (~32 days, term ~67 days), sows were anesthetized in an anesthetic chamber (4-5% isoflurane with 2L/min O_2_, followed by 2.5-3% isoflurane with 1L/min O_2_ for maintenance). Immediately following induction of anesthesia, a subcutaneous injection of Rubinol (Glycopyrrolate, 0.01mg/kg, Sandoz Can Inc., Montreal, QC) was administered. A midline incision was made below the umbilicus, exposing the bicornate uterus. Arterial vessels feeding one horn of the uterus were identified and every second branch was cauterized using an Aaron 2250 electrosurgical generator (Bovie Medical, Clearwater, FL, USA). Immediately after surgery, a subcutaneous injection of Temgesic (Buprenorphine, 0.025mg/kg, Schering-Plough Co., Kenilworth, NJ) was administered, and recorded monitoring continued following surgery for 3 days.

Sows delivered spontaneously at term, at which time pup weight, length, abdominal circumference and biparietal distance were measured. Following the pupping period, all birth weights were collated and if birth weight was below the 25^th^ percentile these guinea pigs were allocated to the low birth weight (LBW) group, and if birth weight was between the 25^th^ and 75^th^ percentile these pups were allocated to the normal birth weight (NBW) group (**Figure 1**).

**Figure 1.**
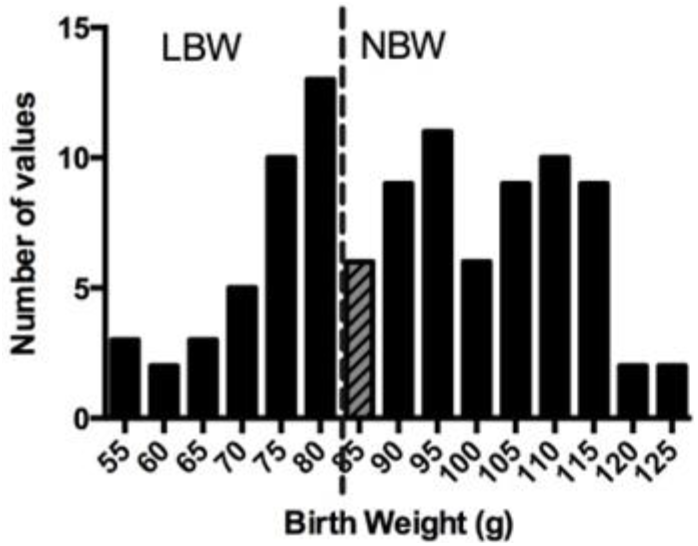
Birth weight distribution. Pregnant guinea pigs underwent uterine artery ablation at mid-gestation. At the end of pupping period, pups born below the 25th percentile (dashed line) were classified as low birth weight (LBW), and those above the 25th percentile were classified as normal birth weight (NBW). Animals between 85g and 90g (grey, hatched bar) were excluded from further analyses.

Based on these criteria, FGR pups were below 80g, and NBW pups were above 90g, in accordance with previously reported pup weight ranges in uterine artery ligation and maternal feed restriction guinea pig models [44]. Animals between 80g and 90g (grey, hatched bar) were excluded from analyses (**Figure 1**).

Pups remained with their dams during the 15-day lactation period. Five days prior to weaning, all pups were introduced to a synthetic control diet (CD, Harlan Laboratories TD.110240) [45] and weaned on postnatal day (PND) 15 into individual cages. At this time, guinea pig offspring were randomized to either a CD or a Western (WD, Harlan Laboratories TD.110239) diet [45]. To avoid litter effects, only one LBW and one NBW animal from a single litter was assigned to each diet. Only male pups were examined for this study. From the time of weaning, food intake was measured daily, and body weights were measured daily until PND50, then twice weekly until PND145, corresponding to young adulthood [46,47].

### 2.3. Analysis of growth and food intake

Absolute growth rate (AGR: g/day), percent increase in body weight, and fractional growth rate (FGR: AGR/birth weight) [46] were calculated for the 15-day lactation period for NBW and LBW offspring.

The post-weaning period was divided into three intervals: weaning-PND50 representing the period of maximal growth, PND51-110 representing the period of adolescent growth, and PND111-145 representing the plateau period of young adulthood. For each interval, the AGR, average food intake (g/day/kg body weight), calorie consumption (calories/day) and efficiency (g gained/day/calories consumed/day) were calculated for NBW/CD, LBW/CD, NBW/WD and LBW/WD offspring.

### 2.4. Assessment of whole-body glucose tolerance

At PND60 and PND120, intraperitoneal glucose tolerance tests (IPGTTs) were performed to assess whole-body glucose tolerance. Following an overnight (16h) fast, a bolus injection of 50% dextrose (1g/kg body weight) was administered intraperitoneally. Blood was sampled at the peripheral ear vein at times 0-, 10-, 20-, 50-, 80-, 110-, 140-, 170-, and 200-minutes following injection. Blood glucose levels were measured using a Bayer Contour hand-held glucometer (Bayer Diabetes, Mississauga, ON). Area under the glucose curve was calculated using GraphPad Prism 6 (GraphPad Software, San Diego, CA) for each animal as a measure of glucose clearance [48].

### 2.5. Assessment of skeletal muscle glucose uptake

Approximately 10 days prior to glucose tolerance testing, at PND50 and PND110, guinea pigs underwent positron-emission tomography (PET) scans in order to assess *in vivo* glucose uptake at the level of the skeletal muscle. Guinea pigs were anesthetized in an anesthetic chamber (4-5% isoflurane with 2L/min O_2_) then transferred to a tight-fitting nose cone (2.5-3% isoflurane with 1L/min O_2_ for maintenance). While anesthetized, each guinea pig was injected with ~18.5MBq of ^18^F-fluoro-deoxy-glucose (FDG) via the pedal vein. Following injection, guinea pigs were recovered and returned to their cages. After an approximately 40 minute period for the uptake of the radiolabeled FDG, guinea pigs were anesthetized (4-5% isoflurane with 2L/min O_2_) and placed on a micro-PET scanner (GE eXplore Vista, GE Healthcare) fitted with a nose cone to inhale 2.5-3% isoflurane with 1L/min O_2_ for maintenance of anesthesia. Emission scans for the leg regions were acquired for 20 minutes, then reconstructed in 3D mode. Resulting images were analyzed with a custom MATLAB-designed program (MATLAB; MathWorks Inc., MA, USA), where regions of interest were drawn around the muscles of the hind-limb. Measurements of Standard Uptake Value (SUV) were obtained by applying FDG dosage and body weight of the pup to the following equation [49,50]:

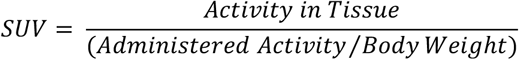

Whole-body adipose tissue, muscle and bone were quantified with computed tomography (CT) at PND110 as previously described [51].

### 2.6. Tissue collection

At PND145, guinea pigs were sacrificed by CO_2_ inhalation [47] following an overnight fast. Gastrocnemius muscle, a representative muscle with mixed composition of oxidative and glycolytic fibers, [52] was removed and trimmed of excess fat and connective tissue, then flash frozen in liquid nitrogen for subsequent analysis. The subset of muscle samples used for histological analysis were fixed in 4% paraformaldehyde and further embedded in paraffin wax (both left and right muscles in a single block).

### 2.7. Determination of fatty infiltration, arteriole density and interstitial fibrosis in skeletal muscle

All tissue processing and cutting, as well as hematoxylin and eosin (H&E) staining were performed at the Molecular Pathology Core Facility (Robarts Research Institute, London, ON, Canada). Cross sections (5μm) were cut towards the middle of the muscle sample. For analysis of muscle fat deposition, sections were stained with H&E. Interstitial adipocytes were not observed within the muscle sections; therefore, fat deposition was not quantified.

To determine arteriole densities, sections were immunostained with alpha-smooth muscle actin (α-SMA) antibody (Clone 1A4, DAKO, M0851, Agilent Technologies, Santa Clara, CA, USA). Slides containing gastrocnemius muscle sections were deparaffinized in xylene and rehydrated using a graded ethanol series. Heat-induced epitope retrieval (HIER) was then performed within an antigen retriever (EMS, Hatfield, PA, USA) using 10mM Tris HCl/ 1mM EDTA pH 9.0. Slides were washed three times in phosphate-buffered saline (PBS). A hydrophobic pen (Dako, Agilent Technologies, Santa Clara, CA) was then used to circle the tissue samples and subsequent incubations were done within a humidified chamber. To reduce non-specific binding, tissues were treated with Background Sniper (Biocare Medical, Concord, CA) for ten minutes. After a brief wash with PBS, primary antibody α-SMA (1:2000 dilution) was applied to the tissue and microscope slides were incubated overnight at 4°C. The following day, slides were rinsed three times with PBS, and the secondary antibody, anti-mouse Alexa 660 (ThermoFisher Scientific, London, ON, Canada) (pseudo-green, 1:400 dilution) was applied. The slides were allowed to incubate for 45 minutes at room temperature, followed by a PBS wash (2×5min). Tissue was then counterstained with 4’, 6-Diamidino-2-Phenylindole, Dihydrochloride (DAPI) (ThermoFisher Scientific, London, ON, Canada) (1:300 dilution) for two minutes, followed by a brief PBS wash. Coverslips were mounted onto the slides (Fisher Scientific, Ottawa, ON, Canada) using 60 μl of Prolong Diamond Mounting Media (ThermoFisher Scientific, London, ON, Canada) and dried overnight prior to imaging.

Imaging and arteriole quantification were performed in a blinded manner. A total of six images spaced across the entirety of the specimen were taken per section using AxioImager Z1 fluorescent microscope (Carl Zeiss Ltd., North York, ON, Canada). Only cross-sectional areas of the muscle were included for quantification. Multichannel images of each field of view included a matching channel image of the natural tissue autofluorescence, for use in measuring total myofiber area. Vessels positive for α-SMA were manually counted, with the exception of larger arteries. Total myofiber areas (mm^2^) were quantified using a customized, automated ImagePro software macro (Media Cybernetics, Rockville, MD). All analysis was done on raw camera files. Binary thresholds for automatically counting myofiber area were generated using the OTSU minimum variance algorithm on the raw greyscale versions of the auto-fluorescent signal channel. Vessel number was then normalized to myofiber area and the resulting densities were averaged across the six images for each guinea pig.

To determine collagen deposition (fibrosis), Masson’s trichrome stain was used. Slides containing muscle sections were deparaffinized in xylene and rehydrated using a graded ethanol series. Tissue was then placed in Harris Hematoxylin solution for 30 seconds, briefly sprayed with water, then dipped three times in HCl (1%) made in ethanol (70%). Slides were then placed in Biebrich Scarlet (Ponceau BS)/Acid Fuchsin (1% Acid Fuchsin/1% Acetic acid) mixture for 2 minutes, followed by a quick rinse in distilled water. Muscle sections were then placed in a solution containing equal parts of phosphotungstic (5%) and phosphomolybdic (5%) acids for 1 minute, followed by 3 min in Fast Green FCF (1%) made in acetic acid (1%). Subsequently, the slides were placed in acetic acid (1%) until collagen retained only a green colour. At this time, tissue was quickly rinsed in 95% alcohol, then dehydrated using a graded series of ethanol, followed by xylene. Coverslips were mounted onto the slides (Fisher Scientific, Ottawa, ON, Canada) using Permount (Fisher Scientific, Ottawa, ON, Canada) and dried overnight before imaging.

Following staining, virtual slides of each specimen were created using an Aperio AT2 Scanner system (Leica Biosystems, Inc., Buffalo Grove, IL, Aperio). Aperio ImageScope v.6.25 software was then used to view the digital slide and extract five images (1890 μm^2^ each) covering the span of the muscle in a blinded manner. Muscle fibrosis areas for each image were determined using a customized, automated ImagePro software macro. Binary counting thresholds for the macro were created through RGB colour samples across a variety of images and tested for accuracy against negative areas of the tissue. Functional collagen surrounding the muscle was excluded from quantification. The collagen areas were normalized to total myofiber area, then averaged across the five images per animal and reported as percentages.

### 2.8. RNA extraction and quantitative real-time PCR (qRT-PCR)

Total RNA was extracted from gastrocnemius muscle by homogenizing ~20mg in 1mL Trizol (Invitrogen, Carlsbad, CA) and following the manufacturers protocol. Total RNA was subsequently treated with deoxyribonuclease for 30 minutes to eliminate genomic DNA. Quantification of RNA was performed with the ND-1000 NanoDrop spectrophotometer (Thermo Fisher Scientific, Rochester, NY). Each sample was also assessed for RNA integrity using 1.2% agarose electrophoresis with RedSafe™ (iNtRON Biotechnology, Sangdaewon-Dong, Joongwon-Ku, Sungnam, Kyungki-Do, 462-120, KOREA). Complementary DNA was synthesized from 2μg of purified RNA, reverse-transcribed using M-MLV Reverse Transcriptase (Life Technologies, Burlington, ON) and a C1000 Thermal Cycler (Bio-Rad, Mississauga, ON). All primers were designed from guinea pig (*cavia porcellus*) sequences using NCBI/Primer-BLAST tool (**Supplementary Table A1**). Standard curves for each primer pair (three pairs for each gene) were generated from serial dilutions of cDNA for determination of primer efficiencies. PCR efficiencies for each primer set were 90-105% (**Supplementary Table A1**). Melting curve analysis and presence of a single amplicon at the expected size in a 1.8% agarose gel were used to confirm amplification of a single product. cDNA products were used as templates for qRT-PCR assessment of gene expression using the SYBR green system (SensiFast^™^ Sybr No-Rox Mix; Bioline, London, UK) on a Bio-Rad CFX384 Real-Time system instrument (Bio-Rad, Mississauga, ON). Each sample was run in triplicate. Forty cycles of amplification were performed, with each cycle consisting of: denaturation at 95°C for 15s, annealing at the pre-determined temperature for each primer set for 20s, and extension at 72°C for 20s. Control samples containing no cDNA were used to confirm absence of DNA contamination. The transcript level of target genes was normalized to β-actin; there were no differences in the expression of β-actin between experimental groups. The fold expression of each individual target gene was determined by the 2^-ΔΔCt^ method [53].

### 2.9. Protein extraction and immunoblotting

Gastrocnemius samples (~50mg each) were homogenized in 0.5mL ice-cold lysis buffer (pH 7.4, 50mM Tris-HCl, 1% NP-40, 0.5% Sodium-deoxycholate, 150mM NaCl, 0.1% sodium dodecyl sulfate) containing protease inhibitor cocktail (Cocktail set 1, Calbiochem, Bilerica, MA) and phosphatase inhibitors (25 mM NaF, 1 mM Na2VO4). Homogenates were incubated on ice for 15 minutes, then subjected to sonication with 3-5 bursts at 30% output (MISONIX: Ultrasonic liquid processor), followed by centrifugation at 15,000g for 30 min at 4°C. The supernatant was collected, and Pierce BCA Protein Assay (Thermo Scientific, Waltham MA) was employed for protein quantification.

Protein samples were separated using 7.5%, 10% or 12% Bis-Tris gels, and transferred onto Immobilon transfer membranes (EMD Millipore, Billerica, MA) via wet transfer. Consistent protein loading and sufficient transfer of proteins was confirmed using 1× Amido Black stain and images were captured using a Versadoc System (Bio-Rad, Mississauga, ON). Membranes were blocked overnight at 4°C with Tris-buffered saline (TBS)/0.1% Tween 20 (Thermo Fisher Scientific, Waltham, MA) containing 5% skim milk or 5% bovine serum albumin (BSA; AMRESCO, Solon, Ohio) as indicated in **Supplementary Table A2**. Following blocking, membranes were incubated with primary antibodies in Tris-buffered saline (TBS)/0.1% Tween 20 containing 5% skim milk or BSA, at 4°C and following dilutions specified in **Supplementary Table A2**. The blots were then washed in TBS/0.1% Tween 20 and incubated at 1h room temperature with the appropriate secondary conjugated antibody (**Supplementary Table A2**). The chemiluminescence signal was captured with the ChemiDoc MP Imaging System (Bio-Rad), and protein band densitometry was determined using the Image Lab software (Bio-Rad). Ponceau staining was used as the loading control, as previously published [54,55]

### 2.10 Thin liquid chromatography and gas chromatography

Total lipids were extracted from gastrocnemius muscle (~160mg), using a protocol adapted from Klaiman et al. [56]. Briefly, tissue samples were homogenized in 850μl 1:1 chloroform:methanol (v/v) containing 0.1% butylated hydroxytoluene (BHT) [57]. Following homogenization, 8.5mL of 2:1 chloroform: methanol (v/v)+0.1% BHT was added to each sample. 1.7mL of 0.25% KCl was added to separate the aqueous solutes, then incubated at 70°C for 10 minutes.

Once the samples were cooled, they were centrifuged at 2000rpm for 5 minutes, and the aqueous layer was removed. The remaining solution, containing lipids, was dried under a gentle stream of N_2_, and resuspended in 1mL chloroform. Twenty μL of total lipids were removed for subsequent analyses and diluted to a concentration of 2μg/μL. The remaining sample was dried again under a gentle stream of N_2_.

Separation and analysis of lipid classes was accomplished by thin layer chromatography-flame ionization detection (TLC-FID) using the Iatroscan MK-6 TLC/FID Analyzer System (Shell-USA, Federicksburg, VA). The system is composed of 10 quartz rods that are 15cm in length and 0.9mm in diameter, coated with a 75μm layer of porous silicagel particulates that are 5μm in diameter (Shell-USA, Federicksburg, VA) [58]. Prior to application of samples, chromarods were repeatedly blank scanned in order to burn off any impurities and activate the rods. 1μl of the 2μg/μL lipid sample or the reference standard was spotted onto the rods. Rods were developed in a TLC chamber with benzene:chloroform:formic acid (70:30:0.5 v/v/v) to separate the neutral lipids until the solvent front reached the 100 mark on the rod holder. Rods were subsequently removed and dried at 60°C in a rod dryer (TK8, Iatron Inc, Tokyo, Japan) for 5 minutes. Rods were analyzed using the following settings: 2L/min atmospheric air, 160mL/min hydrogen, scanning speed of 3s/cm. Area under the curve for the peaks of interest were integrated using PeakSimple 3.29 Software (SRI Instruments, CA) and expressed as a percentage of the total area under the curve for all peaks analyzed.

In preparation for gas chromatography, the total lipids were fractioned using Supelco Supelclear LC-NH_2_ sPE columns (Supelco, Bellefonte, PA). Total lipid samples were resuspended in chloroform to a concentration of 10mg/mL. Once the columns were conditioned, 300μL of suspended sample was added and allowed to flow through the column, then centrifuged at 1000rpm for 1min. Neutral lipids were eluted with 1.8mL of 2:1 chloroform:isopropanol (v/v), followed by centrifugation at 1000rpm for 1min. Non-esterified fatty acids were eluted with 1.6mL 98:2 isopropyl ether:acetic acid (v/v), followed by centrifugation at 1000rpm. Phospholipids were eluted with 3mL of methanol, followed by centrifugation at 1000rpm for 1min. Each sample was subsequently dried under a gentle flow of N_2_. Three hundred μg of each sample was then methylated by adding 200μL of Meth-HCl to the dried sample and incubated at 90°C for 45min. Next, 800μL of water, followed by 500μL of hexane was then added to each sample, with the hexane layer extracted and put in a fresh tube. Following hexane extraction, samples were dried under a gentle flow of N_2_, and resuspended in dichloromethane for injection into a 6890N gas chromatograph (Hewlett Packard, Palo Alto, CA, USA) with a J&W Scientific High Resolution Gas Chromatography Column (DB-23, Agilent Technologies) and a flame ionization detector. Fatty acids were identified by comparison of the relative retention time with known standards (Supelco 37 component FAME mix, and Supelco PUFA No.3, from Menhaden Oil).

### 2.11 Extraction and LC-MS/MS method for the analysis of acylcarnitines and amino acids

Acylcarnitine and amino acid analysis was performed by the Analytical Facility for Bioactive Molecules, The Hospital for Sick Children, Toronto, Canada. Ground tissue was weighed out and a solution of 1:1 water:methanol was added to yield 25mg/mL. Samples were then homogenized and kept on ice. An equivalent of 5mg of homogenized tissue was added to an Eppendorf tube with an acylcarnitine internal standard (IS) mixture (NSK-B, Cambridge Isotopes) and an amino acid IS mixture (Arginine-^13^C_6_, ADMA-d_7_, citrilline-d_7_, glutamic acid-d_2_, ornithine-d_7_, and leucine-d_10_, Cambridge Isotopes and CDN Isotopes). Samples and standard mixtures were then acidified with 60μL of 0.1% formic acid and then protein was precipitated using 1mL of 0.3% formic acid in acetonitrile. Tubes were then vortexed and centrifuged at 10,000g for 10 minutes at 4°C. Supernatants were transferred to conical glass tubes and the remaining pellet in the Eppendorf was re-suspended in an additional 1mL 0.3% formic acid in acetonitrile and vortexed and centrifuged again. The combined supernatants were dried under a gentle flow nitrogen gas. Samples and standards were derivatized with 100μL butanolic-HCL 3N for 20 minutes at 65°C. Solvent was then removed under a gentle flow nitrogen gas and samples were reconstituted in 200μL MPB (see below) and transferred to auto sampler vials. Ten μL of the reconstituted sample were diluted 50x in the same solution for amino acid analysis.

Extracted samples were injected onto a Kinetex HILIC 50 x 4.6 mm 2.6 μm column (Phenomenex) connected to an Agilent 1290 HPLC system attached to a Q-Trap 5500 mass spectrometer (AB Sciex, Framingham, MA). Injected samples were eluted with a gradient of mobile phases MPA: 90/10 5 mM ammonium formate pH 3.2/acetonitrile and MPB: 10/90 5mM ammonium formate pH 3.2/acetonitrile. For Acylcarnitines, the gradient started at 4% MPA for 1 minute, increasing to 45% for 0.65 min, holding for 0.05 min and then returning to 4% MPA for the remaining 3 minutes. For amino acids, the 50 x diluted samples were eluted with a gradient of MPA starting at 4% for 2.5 minutes and increasing to 75% MPA until 6 minutes, holding at 75% MPA for 1 minute, and then returning to 4% over the final 2.5 minutes. Mass transitions were monitored for each sample and compared against known standard mass transitions for species analyzed [59]. Data were collected and analyzed using Analyst v1.6 (AB Sciex, Framingham, MA). Qualitative area ratios were determined for each species examined [59].

### 2.12 Statistical analyses

A Shapiro-Wilks test was used to determine if data was normally distributed, then data were Box-Cox-transformed if necessary, prior to ANOVA. Two-way ANOVA was employed to assess the main effect and the interaction effect of birth weight and postnatal diet (GraphPad Software, San Diego, CA). A Bonferroni post-hoc test was used to compare NBW/CD *versus* LBW/CD as well as NBW/WD *versus* LBW/WD. When comparing only NBW and LBW at birth and during lactation period, two-tailed unpaired *Student’s* t-test was used.

A principal component analysis (PCA) of the acylcarnitine and amino acid measurements was performed (SPSS Statistics, version 21.0; IBM SPSS, Armonk, NY) to reduce the dimensionality of the data set while retaining as much of the variance as possible. The metabolic principal components (PCs), defined as linear, orthogonal (uncorrelated) combinations of the original metabolites, were ordered according to their decreasing ability to explain variance in the original data set. Only PCs with eigenvalues greater than 1 were retained. The cumulative variance explained by the 9 PCs included in the resultant analysis was 87.6%. The data set was then rotated using an orthogonal varimax procedure. To facilitate biological interpretation of the PCs, the associated weight loading factors for each metabolite were examined and metabolites with a loading score greater than 0.5 are reported for each factor. Tests of main effects of birth weight, diet, and birth weight by diet interactions were performed using two-way ANOVA.

Data presented are min. to max. box and whisker plots or mean ± standard error (SEM). Values of p < 0.05 were considered statistically significant for all analyses.

## 3. Results

### 3.1. Characteristics at birth and growth performance during the lactation period

At the time of birth, LBW offspring displayed indications of growth restriction. Birth weight/birth length, a measure of leanness, was significantly (p<0.001) reduced in LBW offspring, compared to NBW offspring (**Table 1**).

**Table 1.**
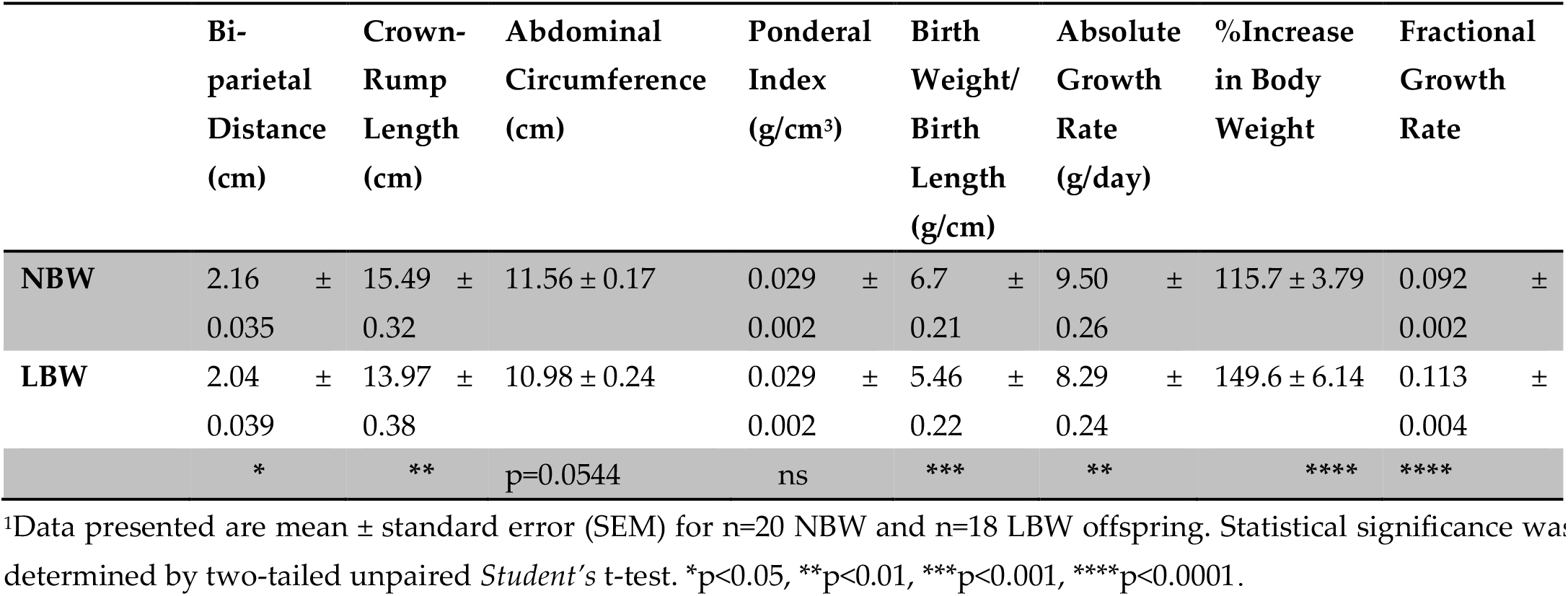
Birth characteristics and lactation period growth.

Biparietal distance (BPD), crown-rump length (CRL) and abdominal circumference (AC) were reduced (p ≤ 0.05) in LBW versus NBW offspring (**Table 1**). During the lactation period, LBW offspring exhibited an altered growth trajectory. LBW offspring showed a 149.6% increase in their body weight over the course of the lactation period, a 29% (p < 0.0001) greater increase than the NBW offspring (**Table 1**). The fractional growth rate was also 23% greater (p < 0.0001) in LBW offspring, compared to NBW (**Table 1**). However, absolute growth rate (AGR) was reduced by 13% (p < 0.01) in LBW offspring (**Table 1**).

### 3.2. Post-weaning growth performance, energy intake and body composition

During the post-weaning period, a difference (p < 0.01) in total mean AGR was also observed in experimental groups, independent of time. Additionally, there was a significant (p < 0.01) interaction between time and our experimental groups (**Figure 2A**). NBW/CD offspring had an AGR of 7.38 ± 0.323 g from weaning until PND50, an AGR of 4.62 ± 0.23 g from PND51-110, and an AGR of 2.87 ± 0.47 g from PND111-145 (**Figure 2A**), ultimately reaching a body weight of 825.2 ± 28.3 g at the time of tissue collection (PND145). During the period from weaning until PND50, NBW/WD offspring displayed a 27% reduction (p < 0.01) in AGR, compared to NBW/CD offspring, whereas LBW/WD offspring displayed a 34% reduction in AGR compared to NBW/CD offspring (p < 0.001; **Figure 2A**). During the same period, no changes in mean AGR was observed in LBW/CD versus NBW/CD. No significant differences in mean AGR were also observed in the 3 groups compared to NBW/CD for the period from PND51-110. From PND111 to the time of euthanasia, LBW/CD offspring displayed a 52% reduction (p < 0.05) in AGR, compared to NBW/CD offspring (**Figure 2A**). Overall, WD-fed offspring remained significantly (p < 0.05) lighter than CD-fed offspring at the time of tissue collection, with body weights of NBW/WD: 732.3 ± 26.8 g; LBW/WD: 708.8 ± 19.8 g vs NBW/CD: 825.2 ± 28.3 g; LBW/CD: 747.9 ± 38.1 g. Compared to the NBW/CD group, LBW/WD animals had significantly (p < 0.05) lower body weights at euthanasia.

**Figure 2.**
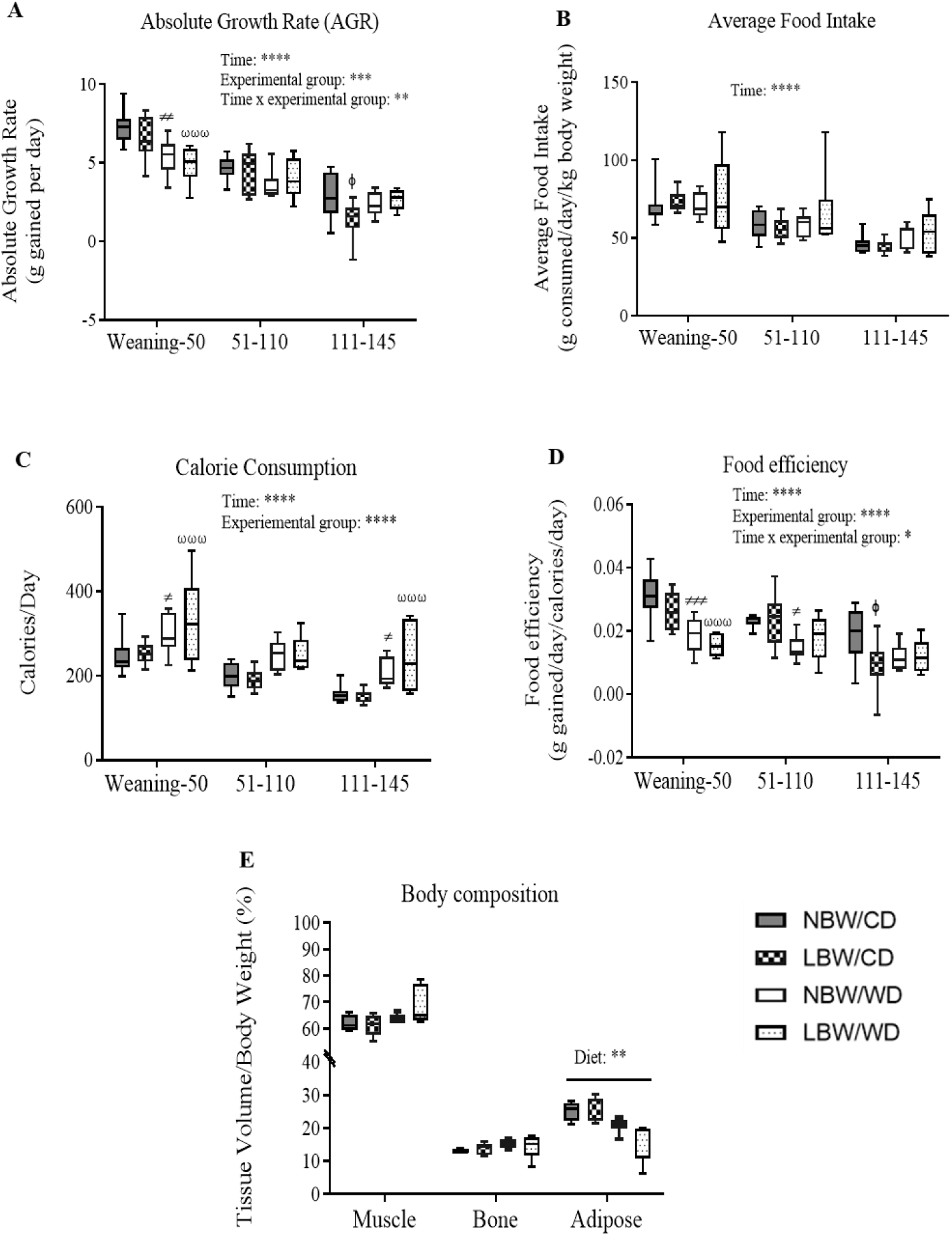
Post-weaning growth and feeding behavior. **A**) Absolute growth rate (AGR) = grams gained per day. **B**) Average food intake = grams of food consumed/kilogram body weight. **C**) Calorie consumption = calories consumed per day. **D**) Food efficiency = grams gained per day/calories consumed per day. Data presented are min. to max. box and whisker plots for 6-11 animals/experimental group. Two-way repeated-measures ANOVA showed that the main effects of experimental group, time as well as the time × experimental group interaction: **p < 0.01; ***p < 0.001; ****p < 0.0001. ^ϕ^p < 0.05 for the difference between NBW/CD and LBW/CD, ≠p < 0.05; ≠≠p < 0.01 for the difference between NBW/CD and NBW/WD, and^ωωω^p < 0.001 for NBW/CD versus LBW/WD following a Bonferroni post hoc test. **E**) For body composition at 120 days, statistical significance was determined by ordinary Two-way ANOVA, followed by Bonferroni post hoc test on 4-5 animals per experimental group. **p < 0.01 for the main effect of diet.

Food intake (g consumed/day/kg body weight) was reduced as a main effect of time but no significant differences in average food intake were observed between the four experimental groups throughout the experiment (**Figure 2B**). However, calorie consumption (calories/day) was overall significantly (p < 0.0001) higher in WD-fed offspring, independent of time (**Figure 2C**). From weaning-PND50, calorie consumption was 23% higher in NBW/WD (p < 0.05) and 36% higher in LBW/WD (p < 0.001) offspring compared to NBW/CD (**Figure 2C**). From PND111-PND145, calorie consumption was higher 34% in NBW/WD (p < 0.05) and 55% higher in LBW/WD (p < 0.001) when compared to NBW/CD (**Figure 2C**).

Overall, food efficiency (grams body weight gained/day/calories consumed/day) was significantly (p<0.0001) decreased in WD-fed offspring (**Figure 2D**). From weaning-PND50, food efficiency was reduced by 39% in NBW/WD (p < 0.001) and by 52% in LBW/WD (p < 0.001) offspring compared to NBW/CD. From PND51-110, NBW/WD offspring showed a 35% reduction in (p < 0.01) food efficiency compared to NBW/CD, whereas LBW/CD offspring showed a 50% reduction in (p < 0.01) food efficiency from PND111-PND145, when compared to NBW/CD (**Figure 2D**).

Whole-body adipose tissue, as measured by CT at PND110, was significantly (p<0.01) reduced in WD-fed offspring, independent of birth weight (**Figure 2E**).

### 3.3. Muscle arteriole density, interstitial fibrosis and fatty infiltration

To determine arteriole densities, sections were stained for α-SMA at PND145 (**Figure 3A**). Vessels positive for α-SMA were manually counted and normalized to total myofiber areas. LBW guinea pigs, irrespective of the consumed postnatal diet, displayed a 22% decrease in number of arterioles per myofiber area (p < 0.05, **Figure 3B**). There were no significant differences in arteriole densities between dietary groups.

**Figure 3.**
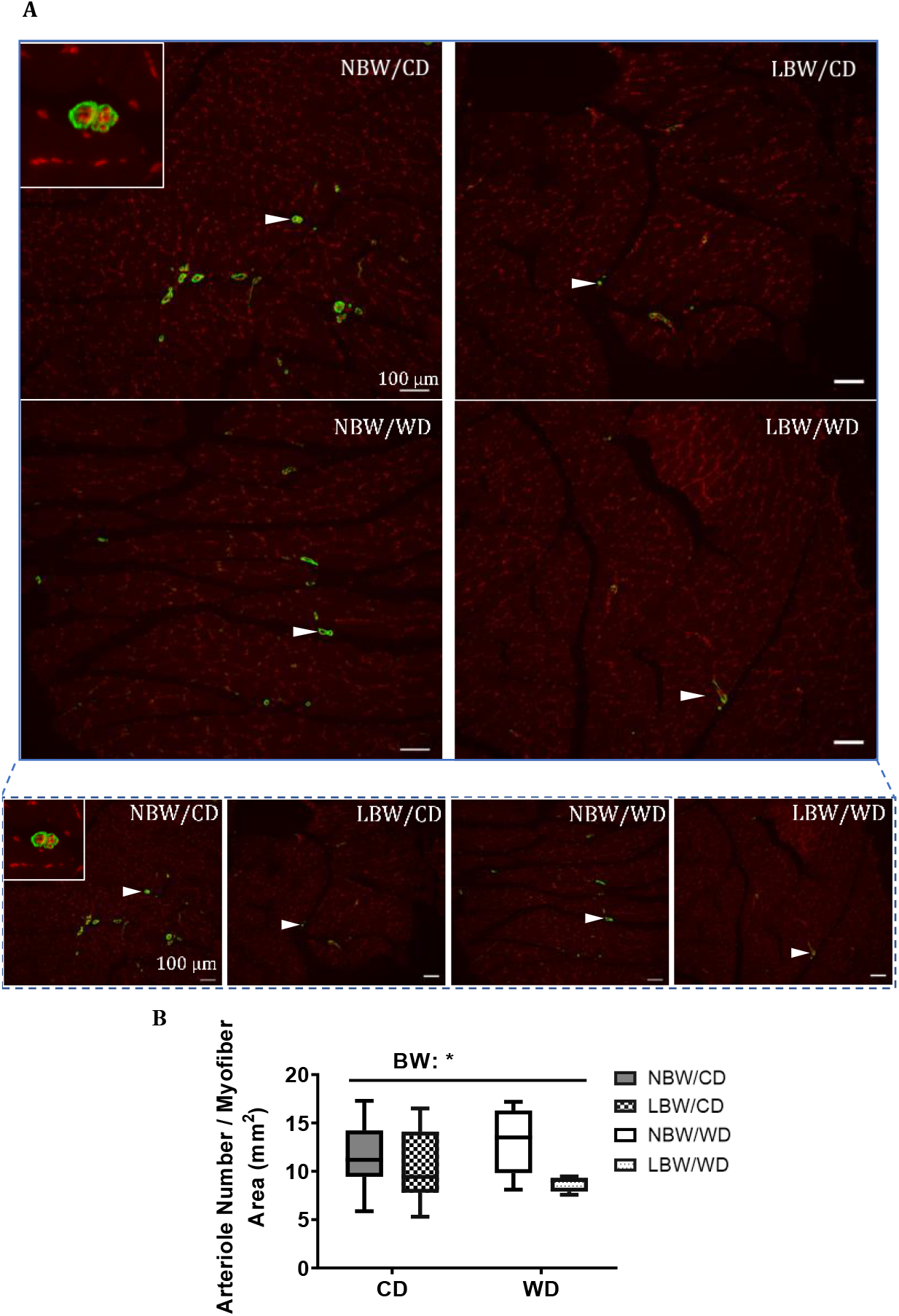
Microtome sections of smooth muscle α-actin (α-SMA) immunoreactivity of muscle arterioles. Sections were immunostained for α-SMA (white arrowhead) to identify vascular smooth muscle cells in gastrocnemius from 145 day-old NBW and LBW guinea pig offspring fed a postnatal CD or WD. (**A**) Representative fields at high and low magnification and (**B**) arteriole number/myofiber area are displayed. Data are presented as min. to max. box and whisker plots for 5-9 animals/experimental group. BW, birth weight; scale bar = 100 μm. *p < 0.05 for the main effect of birth weight (BW).

Collagen deposition between myofiber bundles was determined using Masson’s Trichrome stain at PND145 (**Figure 4A**). All collagen areas were quantified, normalized to total myofiber area, and reported as percentages. A tendency to increased interstitial collagen deposition per myofiber area in LBW offspring, irrespective of diet, was observed (p = 0.064, **Figure 4B**). Consumption of a postnatal Western diet at this age did not have any significant effects on muscle fibrosis.

**Figure 4.**
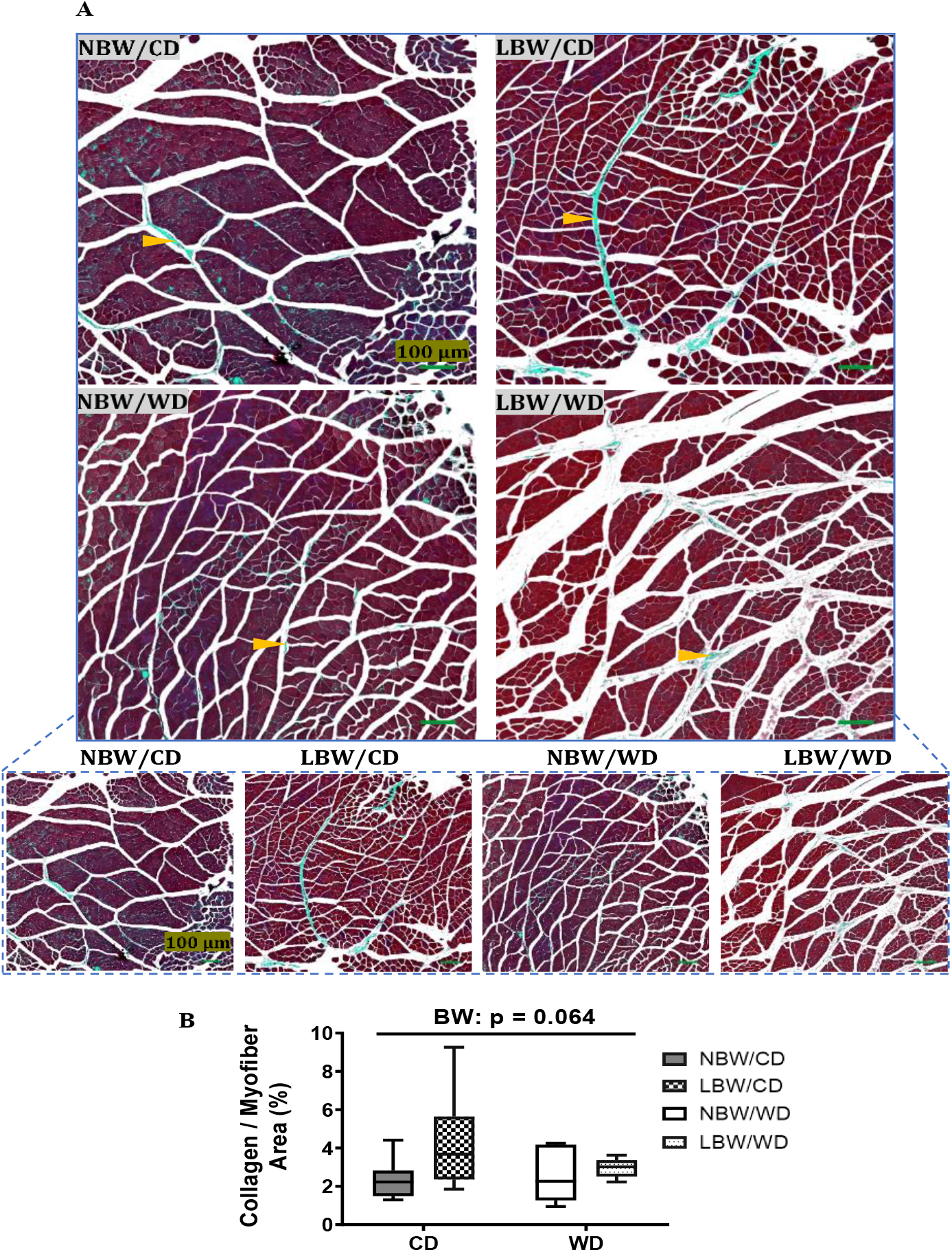
Muscle fibrosis in guinea pig offspring. Sections were stained with Masson’s trichrome to identify collagen deposition (orange arrowheads) at postnatal day 145 in guinea pig offspring fed a postnatal CD or WD. (**A**) Representative fields at high and low magnification and (**B**) percentage of collagen deposition/myofiber area are displayed. Data are presented as min. to max. box and whisker plots for 5-9 animals/experimental group.

The accumulation of adipocytes within muscle is a common manifestation of muscle pathology and was investigated in hematoxylin and eosin (H&E) stained sections of gastrocnemius from NBW and LBW offspring fed the CD or WD. Interstitial adipocytes were not observed in histological images for any treatment groups **(Figure 5**); therefore, fat deposition was not quantified.

**Figure 5.**
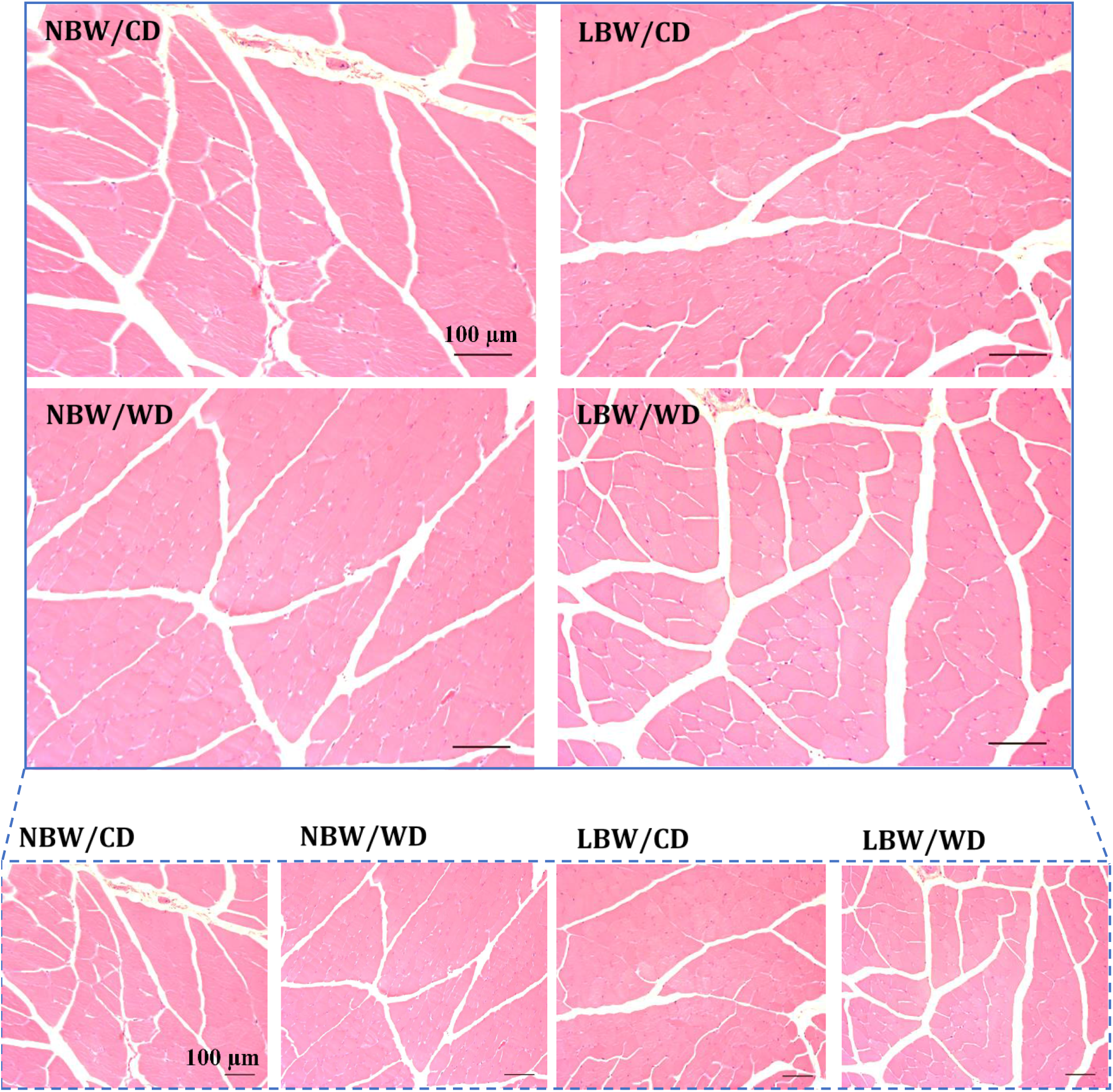
Hematoxylin and eosin (H&E) staining of gastrocnemius muscle. Gastrocnemius was isolated from 145-day-old. Cross sections (5 μm) from 5-9 animals/experimental group were stained with H&E to identify lipid vesicles. Representative fields are shown at low and high magnification are displayed. Interstitial adipocytes were not present for any treatment group, and therefore not quantified. Scale bar = 100 μm.

### 3.4. Whole body and muscle specific glucose homeostasis

Intraperitoneal glucose tolerance challenges were performed to assess whole body glucose tolerance. At PND 60, the blood glucose level reached a peak at 50 min after the glucose injection, and the glucose concentrations were higher in LBW/CD and NBW/WD compared to NBW/CD at 50-, 80- and 110-min post injection (p < 0.05; **Figure 6A**). Furthermore, the whole-body glucose excursion after glucose loading, as measured by the area under the curve, decreased in LBW/WD compared to NBW/WD (p = 0.05, **Figure 6C**). No significant differences were observed in blood glucose levels and whole-body glucose handling following a glucose load at PND120 (**Figure 6B, D**).

Glucose uptake at the level of the skeletal muscle was assessed using ^18^F-FDG PET and measuring the standard uptake value of radioactivity observed during the scans. No significant differences in skeletal muscle glucose uptake were observed due to birth weight or postnatal diet at either PND50 (**Figure 6E**) or PND110 (**Figure 6F**).

**Figure 6.**
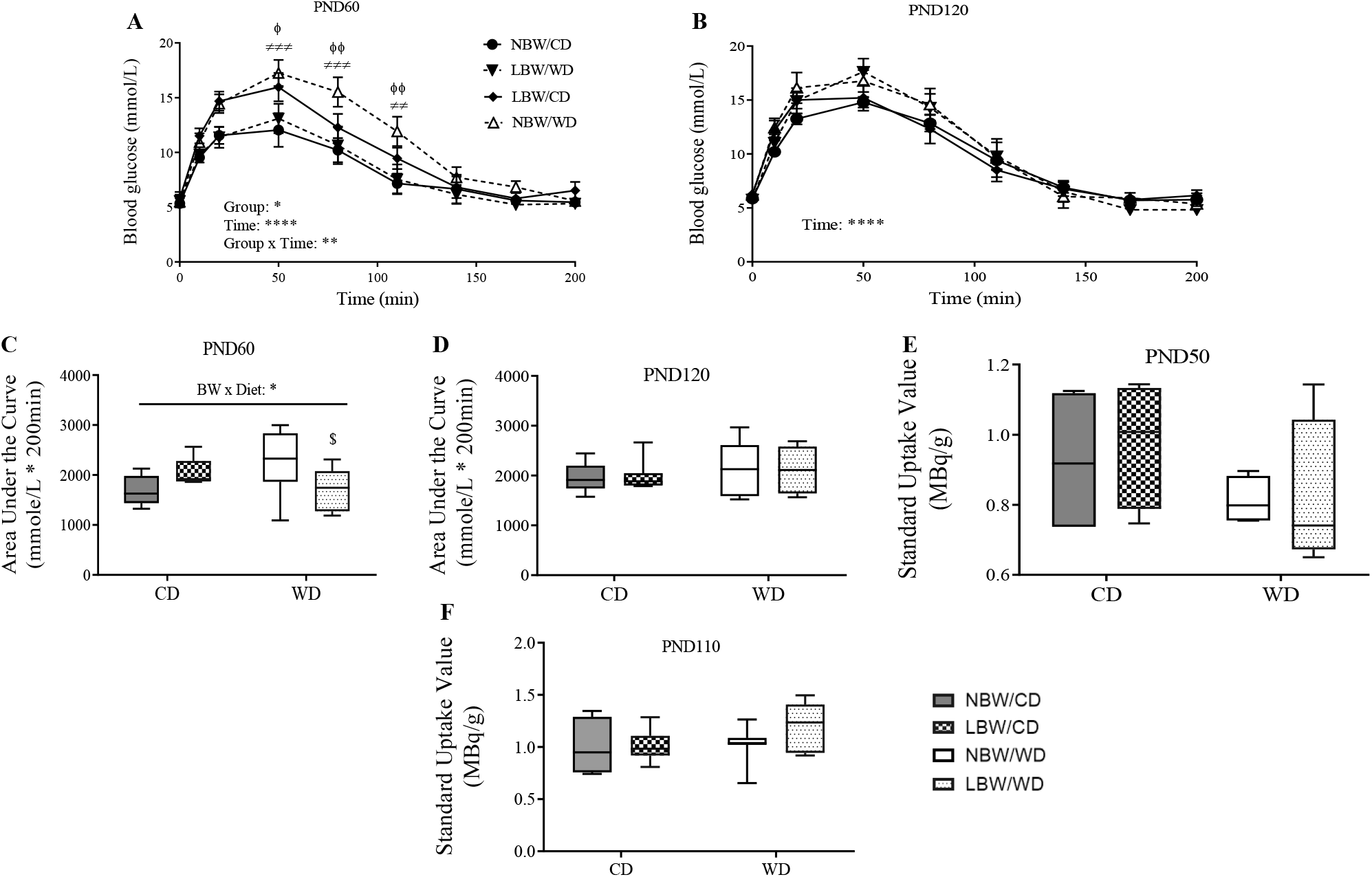
Whole-body glucose tolerance and skeletal muscle glucose uptake. Blood glucose at PND60 (**A**) and PND120 (**B**) was measured after an overnight (16h) fast, and at specific time points for 200 min following an intraperitoneal injection of 1g/kg of 50% dextrose solution. The area under the glucose curve (AUC) as an index of whole-body glucose excursion after glucose loading at postnatal day (PND)60 (**C**) and PND120 (**D**). Standard Uptake Value (SUV) = image derived radioactivity/(injected activity/body weight) at PND50 (**E**) and PND110 (**F**) are presented. Data presented are the mean ± standard error (SEM) or min. to max. box and whisker plots for n = 4-12 animals/experimental group. Twoway repeated-measures ANOVA showed that the main effects of experimental group, time as well as the time × experimental group interaction: *p < 0.05; **p < 0.01; ****p < 0.0001. ^ϕ^p < 0.05 and ^ϕϕ^p < 0.01 for the difference between NBW/CD and LBW/CD and ^≠≠^p < 0.01; ^≠≠≠^p < 0.001 for the difference between NBW/CD and NBW/WD following a Bonferroni post hoc test. BW x Diet: * for p < 0.05 for the main effect of birth weight (BW) and diet interaction using a regular two-way ANOVA.

### 3.5. Skeletal muscle lipid classes and fatty acid profile

Since aberrant lipid storage or lipid intermediates may be involved in diabetes pathogenesis, separation and determination of lipid classes in gastrocnemius muscle was accomplished using thin layer chromatography-flame ionization detection (TLC-FID). At PND150, there were no significant differences in the accumulation of triglycerides (TG), cholesterol (Chol), diacylglycerol (DAG) or phospholipids (PL) due to birth weight or postnatal diet (**Figure 7A**). The fatty acid pool in gastrocnemius was profiled using gas chromatography. WD-fed offspring displayed significantly more myristic acid (C14:0), and less linoleic acid (C18:2n6c) and α-linolenic (C18:3n3) in their neutral lipid fraction, irrespective of birth weight compared to CD-fed offspring (p < 0.01, **Figure 7B**). In the phospholipid fraction, there was a significant decrease in palmitic acid (C16:0) and an increase in octadecenoic acid (C18:1n7) in WD-fed animals (**Figure 7C**). No alterations in the fatty acid composition were influenced by birth weight alone.

**Figure 7.**
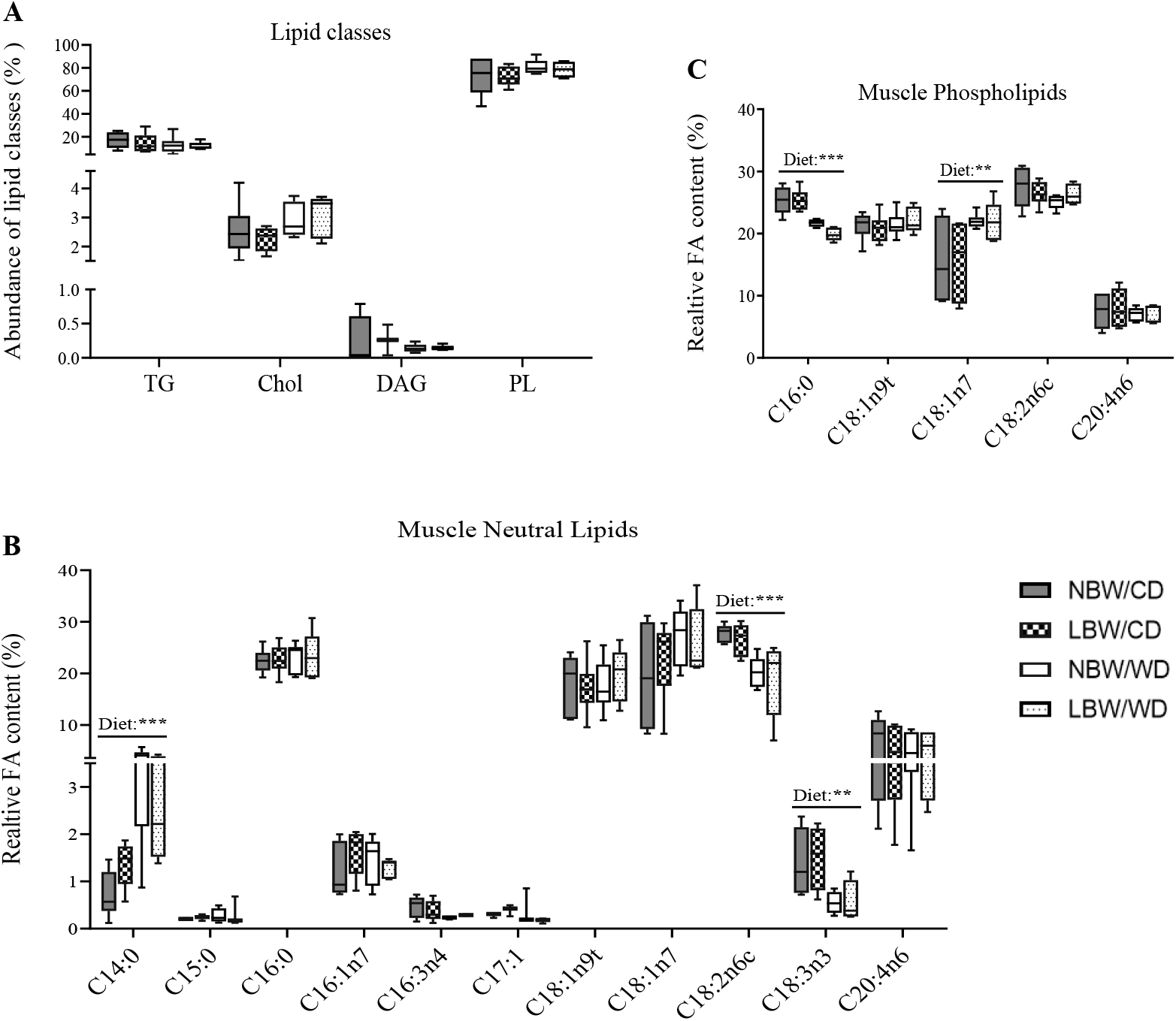
Relative abundance of lipid species and fatty acid profiles at postnatal day 145. (**A**) Relative abundance of lipid species including triglycerides (TG), cholesterol (Chol), diacylglycerols (DAG) and phospholipids (PL) are displayed. Fatty acids in neutral lipid (**B**) and phospholipids (**C**) fractions are displayed. Data are min. to max. box and whisker plots for 4 to 6 animals/experimental group. *, ** and *** indicate *p* < 0.05, *p* < 0.01 and 0.0001 respectively, for the main effect of diet using a regular two-way ANOVA.

### 3.6. Markers of mitochondrial overload and altered amino acid profile in muscle of WD-fed and LBW offspring

Accumulation of even-chain acylcarnitines represents partially oxidized intermediates of β-oxidation, whereas accumulation of odd-chain acylcarnitines represents catabolism of amino acids. In gastrocnemius muscle, medium-chain acylcarnitines (C6-C12) were significantly increased following consumption of a postnatal WD (p < 0.05, **Figure 8A**). Additionally, a significant (p < 0.05) effect of birth weight upon accumulation of C8 and C10 was also observed (**Figure 8A**). Furthermore, an interactive effect of birth weight and postnatal diet was observed in C8, C10 and C12 acylcarnitines, with LBW/WD offspring exhibiting increased accumulation of these acylcarnitines when compared to NBW/WD offspring (p < 0.05, **Figure 8A**). Level of C12 was also increased (p < 0.05) in LBW/CD versus NBW/CD. Levels of C2 acylcarnitine, the end product of complete β-oxidation, were significantly reduced in WD-fed offspring (p < 0.05, **Figure 8A**). Similarly, long-chain acylcarnitines including C14, C14:1, C16, C16:1, C18 and C18:1, were significantly increased following consumption of a postnatal WD (p < 0.001, **Figure 8B**). Significant main effects of birth weight were observed in C14:1, C16, C18 acylcarnitines (**Figure 8B**). A significant interactive effect between birth weight and postnatal diet was observed for C14:1, C16 and C18 acylcarnitines with the magnitude of increase of these acylcarnitines being more elevated in LBW/WD compared to NBW/WD (p < 0.05, **Figure 8B**). A significant reduction in C20:4 acylcarnitine levels was also observed in WD-fed offspring (p < 0.01, **Figure 8B**).

**Figure 8.**
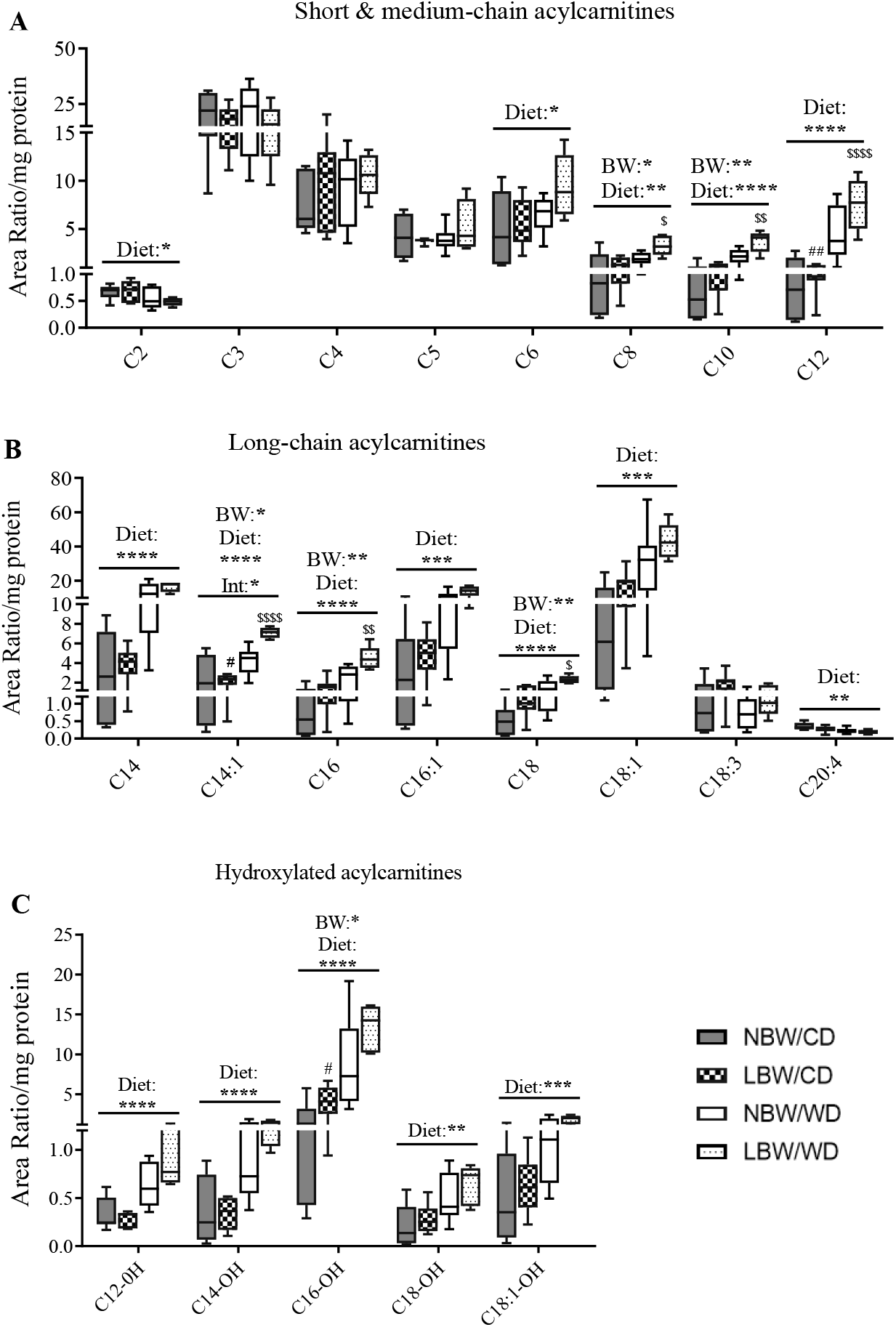
Acylcarnitine contents in offspring’s muscle at postnatal day 145. Short-, medium- (**A**), long- (**B**) chain and hydroxylated (**C**) acylcarnitines are presented. Data are min. to max. box and whisker plots for 5 to 8 animals/experimental group. *, **, *** and **** for p < 0.05, p <0.01, p <.001 and p <.0001, respectively, represent the main effect of birth weight (BW), diet (Diet) or their interaction (BW x Diet) using a regular two-way ANOVA. # and ## indicate p <0.05 and 0.01 respectively, when comparing NBW/CD versus LBW/CD using a Bonferroni post-hoc test. ^$^, ^$$^ and ^$$$$^ denote *p* < 0.05, 0.01 and 0.000 respectively, for NBW/WD versus LBW/WD by Bonferroni post-hoc test.

As inhibition of NADH-linked β-hydroxy fatty acyl-CoA oxidation is known to promote accumulation of hydroxyacylcarnitines, we also sought to quantify these intermediates. We observed significant (p < 0.05) accumulation of the 3-β-hydroxylated forms of C12, C14, C16, C18 and C18:1 in response to WD-feeding (p < 0.01, **Figure 8C**). Birth weight was also associated with a significant accumulation of C16-OH acylcarnitine, with LBW/CD exhibiting more C16-OH than NBW/CD (p < 0.05, **Figure 8C**).

Clusters of amino acids including the branched-chain amino acids (Leu, Iso, Val), as well as Tyr and Phe have recently been identified as strong predictors of increased risk of developing diabetes and impaired insulin action, prompting us to investigate amino acid profiles in muscle of NBW and LBW offspring fed a postnatal CD or WD. In our model, a main effect of birth weight on Glu concentration was observed with LBW/CD displaying less Glu than NBW/CD group (p < 0.05, **Figure 9A**). Although not significant, levels of Leu in LBW offspring, irrespective of postnatal diet tended to increase (p = 0.053, **Figure 9C**). Additionally, we observed increased Ala, Asp and Phe and reduced Cit as main effect of WD irrespective of birth weight (p < 0.05, **Figures 9A, B**).

**Figure 9.**
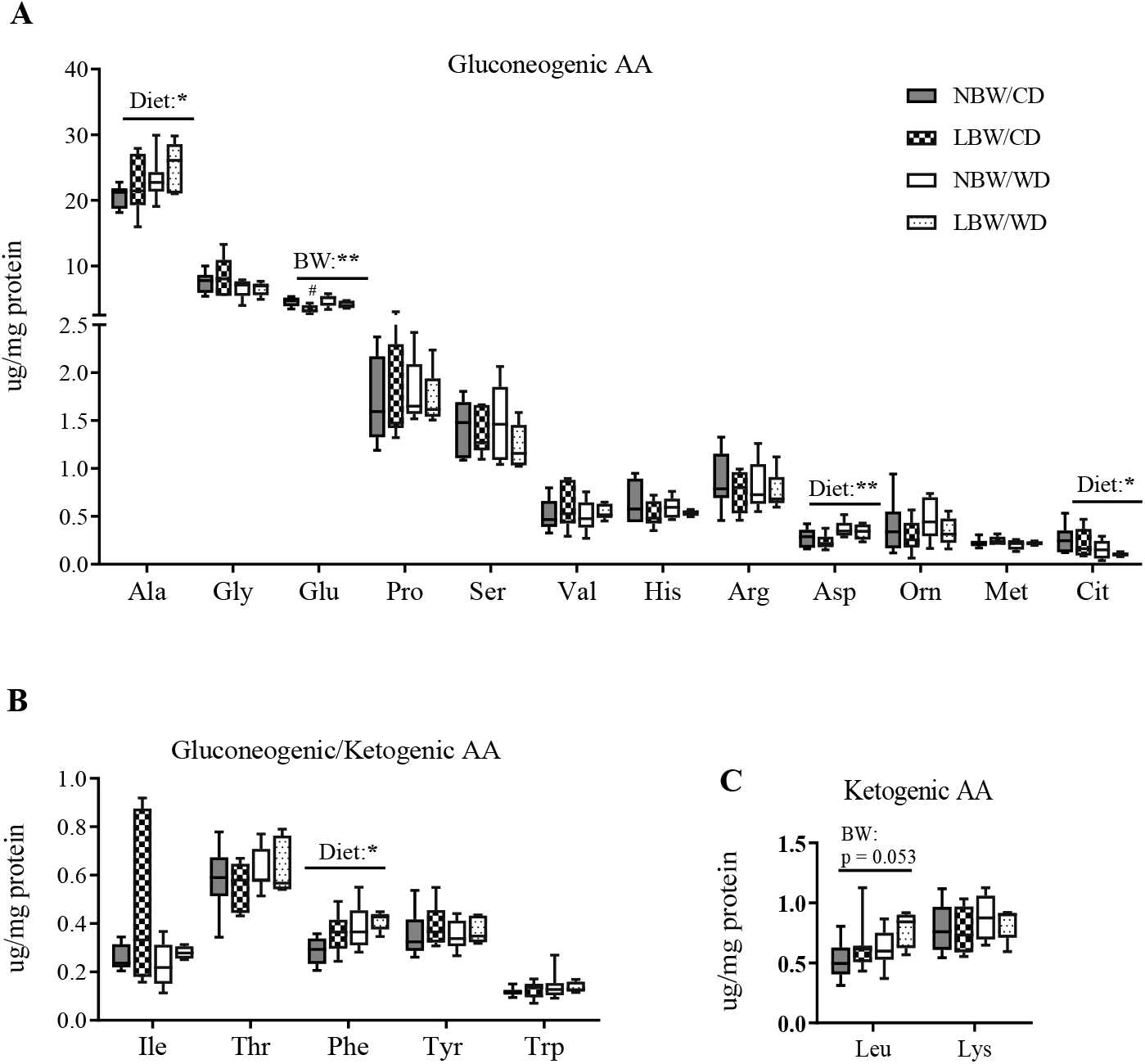
Amino acid (AA) concentrations in offspring’s muscle at postnatal day 145. Gluconeogenic (**A**), gluconeogenic/ketogenic (**B**) and ketogenic AA (**C**) are presented. Data are min. to max. box and whisker plots for 5 to 8 animals/experimental group. * and ** for p < 0.05 and p < 0.01, respectively, represent the main effect of birth weight (BW) or diet (Diet) using a regular two-way ANOVA. # indicates p <0.05 and 0.01 when comparing NBW/CD versus LBW/CD using a Bonferroni post-hoc test.

### 3.7. Principle components analysis (PCA) of muscle acylcarnitines and amino acids

We employed PCA as an unbiased strategy to reduce the dimensionality of acylcarnitines and amino acid data set while retaining as much of the variance as possible. Nine principal components (PCs) were identified, explaining 87.6% of the total variance in the model. Features of the top 4 PCs are described in **Table 2**. The first PC (PC1; medium- and long-chain acylcarnitines) was composed of even-chain acylcarnitines ranging from C8 through C18, in addition to the unsaturated acylcarnitines C14:1, C16:1 and C18:1. Additionally, C2 acylcarnitine was also a part of this component, however the negative loading factor suggests that levels of C2 would decrease as levels of the other components of this cluster increase, similar to the accumulation profile we observed in **Figure 7**. A significant (p < 0.0001) main effect of diet was observed for the components of this factor, highlighting postnatal diet as a major contributor to the acylcarnitine species associated with this component.

**Table 2.**
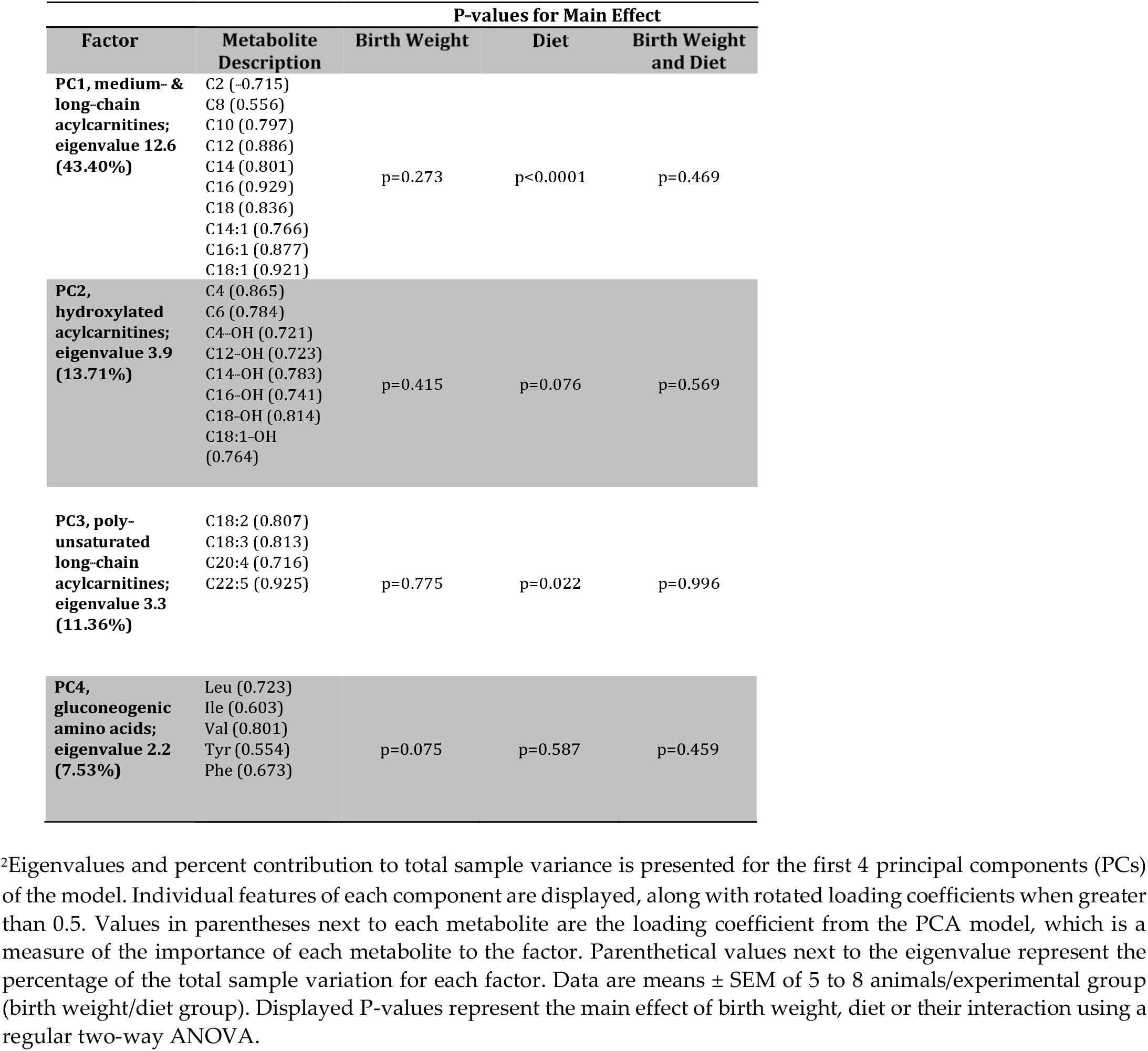
Exploratory PCA on acylcarnitine levels.

| Factor | Metabolite Description | P-values for Main Effect |  |  |
| --- | --- | --- | --- | --- |
|  |  | Birth Weight | Diet | Birth Weight and Diet |
| **PC1, medium- & long-chain acylcarnitines; eigenvalue 12.6 (43.40%)** | C2 (-0.715) |  |  |  |
|  | C8 (0.556) |  |  |  |
|  | C10 (0.797) |  |  |  |
|  | C12 (0.886) |  |  |  |
|  | C14 (0.801) | p=0.273 | p<0.0001 | p=0.469 |
|  | C16 (0.929) |  |  |  |
|  | C18 (0.836) |  |  |  |
|  | C14:1 (0.766) |  |  |  |
|  | C16:1 (0.877) |  |  |  |
|  | C18:1 (0.921) |  |  |  |
| **PC2, hydroxylated acylcarnitines; eigenvalue 3.9 (13.71%)** | C4 (0.865) |  |  |  |
|  | C6 (0.784) |  |  |  |
|  | C4-OH (0.721) | p=0.415 | p=0.076 | p=0.569 |
|  | C12-OH (0.723) |  |  |  |
|  | C14-OH (0.783) |  |  |  |
|  | C16-OH (0.741) |  |  |  |
|  | C18-OH (0.814) |  |  |  |
|  | C18:1-OH (0.764) |  |  |  |
| **PC3, poly-unsaturated long-chain acylcarnitines; eigenvalue 3.3 (11.36%)** | C18:2 (0.807) |  |  |  |
|  | C18:3 (0.813) |  |  |  |
|  | C20:4 (0.716) | p=0.775 | p=0.022 | p=0.996 |
|  | C22:5 (0.925) |  |  |  |
| **PC4, gluconeogenic amino acids; eigenvalue 2.2 (7.53%)** | Leu (0.723) |  |  |  |
|  | Ile (0.603) |  |  |  |
|  | Val (0.801) | p=0.075 | p=0.587 | p=0.459 |
|  | Tyr (0.554) |  |  |  |
|  | Phe (0.673) |  |  |  |

<sup>2</sup>Eigenvalues and percent contribution to total sample variance is presented for the first 4 principal components (PCs)
of the model. Individual features of each component are displayed, along with rotated loading coefficients when greater than 0.5. Values in parentheses next to each metabolite are the loading coefficient from the PCA model, which is a measure of the importance of each metabolite to the factor. Parenthetical values next to the eigenvalue represent the percentage of the total sample variation for each factor. Data are means ± SEM of 5 to 8 animals/experimental group (birth weight/diet group). Displayed P-values represent the main effect of birth weight, diet or their interaction using a regular two-way ANOVA.

PC2 was composed predominantly of the hydroxylated acylcarnitines derived from C4, C12-C18 and C18:1, with contributions from C4 as well as C6. A potential contribution (p = 0.076) of postnatal diet was observed for this PC.

PC3 was made up of poly-unsaturated long-chain acylcarnitines including C18:2, C18:3, C20:4 and C22:5 acylcarnitine species, with a significant (p < 0.05) contribution of postnatal diet to this cluster. PC4 (gluconeogenic amino acids) was composed of the amino acids including Leu, Iso, Val, Tyr and Phe. Birth weight was a potential (p = 0.075) contributor to this PC.

### 3.8. Skeletal muscle insulin signaling signature

The phosphorylation of the β-subunit of the insulin receptor (IRβ) is required for its activation and propagation of downstream signaling events. Phosphorylation of IRβ^Tyr1150/1151^ was not altered by birth weight or diet (**Figure 10**). Protein Kinase B (Akt) is phosphorylated by Phosphoinositide 3-kinase (PI3)-kinase at two resides, Ser^473^ and Thr^308^, that are both required for full activation of this protein. At the Ser^473^ and Thr^308^ residues, phosphorylation was reduced in WD-fed offspring, irrespective of birth weight (p < 0.05 **Figure 10**). Additionally, as a birth weight effect, phosphorylation at Thr^308^ residue was increased (p < 0.05). Stress-induced kinases, including c-Jun N-terminal kinases (JNK), Protein kinase C (PKC)θ, PKCε are known to be activated in situations of oxidative stress or lipid accumulation and insulin resistance. In our study, total protein level of PKCε was increased as a main effect of birth weight with LBW/CD displaying more PKCε level than NBW/CD (p < 0.05 **Figure 10**). Total protein levels of PKCθ and PKCε were also significantly higher in WD-fed animals (p < 0.05, **Figure 10**). JNK1-mediated disruption of insulin receptor/Insulin receptor substrate 1 (IRS1) interaction and the resulting insulin resistance are dependent on IRS1 phosphorylation at both Ser^302^ and Ser^307^. IRS1 is also phosphorylated at Ser^307^ by IKKβ kinase. In the current study, while activation of JNK through phosphorylation at Thr^183^/Tyr^185^ was not altered by birth weight nor diet, phosphorylation of IRS1 at Ser^302^ increased in WD-fed offspring (p <0.0001, **Figure 10**). Interestingly, IRS1 phosphorylation at Ser^302^ was exacerbated in LBW/WD compared to NBW/WD (p <0.001, **Figure 10**). The activation of IKKβ through phosphorylation at Ser^177/181^ was also increased in WD-fed offspring (p <0.0001, **Figure 10**). The first step by which insulin increases energy storage or utilization in muscle involves the regulated transport of glucose into cells, mediated by the facilitative glucose transporter 4 (GLUT4). While GLUT4 protein was not measured, total GLUT5, was significantly (p < 0.0001) increased as a WD main effect (**Figure 10**).

**Figure 10.**
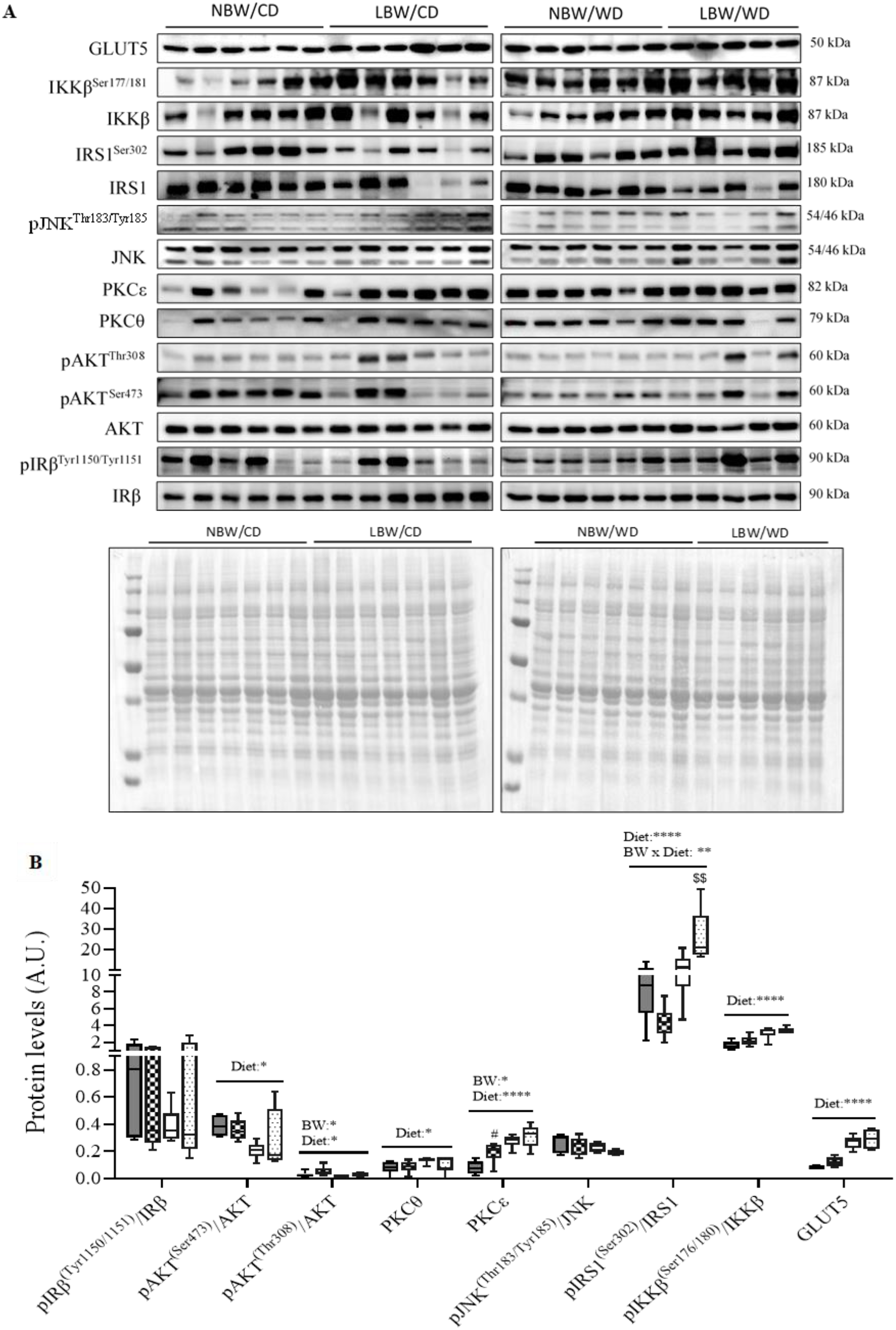
Levels of proteins involved in insulin signaling and glucose transport at postnatal day 145. Panel **A** shows representative cropped blots; panel **B** indicates normalized densitometry values of proteins; Data are min. to max. box and whisker plots for 5 to 6 animals per group. *, ** and ****for p < 0.05, p < 0.01 and p < 0.0001, respectively, represent the main effect of birth weight (BW) or diet (Diet) using a regular two-way ANOVA. ^#^ denotes significant difference (*p* < 0.05) between NBW/CD versus LBW/CD; $$ indicates p <0.01 when comparing NBW/WD versus LBW/WD by Bonferroni post-hoc test.

### 3.9. Expression of genes involved in mitochondrial β-oxidation

The rate limiting step of mitochondrial β-oxidation, CPT1-catalyzed transport of acylcarnitines into the mitochondria, is often used as a marker of mitochondrial oxidative capacity. We observed a significant (p < 0.05) decrease in *CPT1b* mRNA expression in LBW offspring, irrespective of postnatal diet (**Figure 11A**). The 3-ketoacyl-CoA thiolase (KT), the enzyme responsible for catalyzing the removal of acetyl-CoA from the fatty acid chain, was significantly (p < 0.05) decreased at mRNA level in skeletal muscle of LBW offspring, irrespective of postnatal diet (**Figure 11A**).

**Figure 11.**
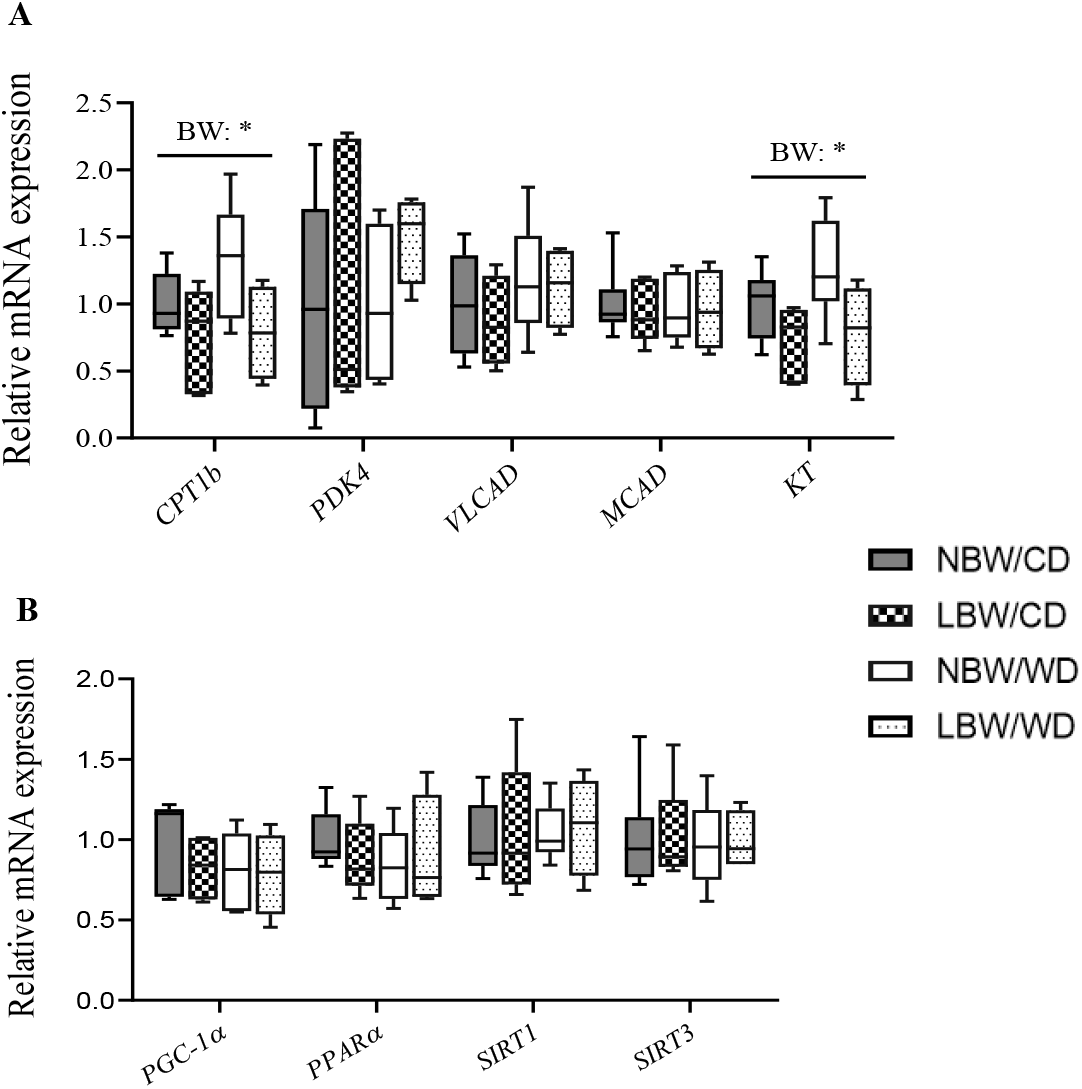
mRNA expression of genes involved in mitochondrial β-oxidation and biogenesis. at postnatal day 145. Panel **A** shows mRNA expression of genes of mitochondrial β-oxidation; panel **B** indicates mRNA expression of transcription factors regulating mitochondrial β-oxidation genes. Data are min. to max. box and whisker plots for n = 4 to 6 animals per group. * for p < 0.05, represent the main effect of birth weight (BW) using a regular two-way ANOVA.

PGC-1α and PPARα are well-known transcription factors that regulate aspects of mitochondrial lipid oxidation. Skeletal muscle *PGC-1α* and *PPARα* mRNA expression was similar among groups (**Figure 11B**). Skeletal muscle sirtuins are a family of transcription factors regulating mitochondrial biogenesis and function. Similar to *SIRT1, SIRT3* mRNA expression was not affected by birth weight or postnatal diet (**Figure 11B**).

## 4. Discussion

Intrauterine growth restriction (IUGR) and low birth weight (LBW) are the results of survival mechanisms adopted by the fetus following uteroplacental insufficiency (UPI), a condition where placenta is unable to deliver an adequate supply of nutrients and oxygen to the fetus. These outcomes have immediate implications for neonatal morbidity and mortality [60,61], but are also now recognized as setting the stage for an increased risk of metabolic 843 dysregulations and metabolic syndrome (MetS) later in life [19,62–64]. Central to MetS is muscle dysfunction and in particular muscle insulin sensitivity and metabolic function [65]. In addition, IUGR resulting in low birth weight (LBW) is now recognized as a central programming factor in lifelong impairments in muscle development and metabolism [24]. With IUGR/LBW outcomes continuing to increase [40] and the increasing availability and consumption of energy-dense diets in society [66], a significant contributor to MetS [67], the present investigation sought to identify alterations in muscle metabolic function. Specifically, the study focused upon changes in skeletal muscle arteriole density, fibrosis, amino acid and mitochondrial lipid metabolism, markers of insulin signaling, and glucose uptake following UPI-induced IUGR/LBW independently, and in conjunction with early postnatal exposure to a WD. The key findings from the present study are that UPI-induced IUGR/LBW and WD affect offspring skeletal muscle amino acid and acylcarnitine profile at early adulthood, and specific changes in muscle acylcarnitines in UPI-induced IUGR/LBW offspring are exacerbated by intake of postnatal WD, in the absence of obesity. Additionally, UPI-induced IUGR/LBW, was associated with notable diminished arteriole density and significant increase in PKCε protein levels in skeletal muscle at early adulthood, in the absence of alterations in skeletal muscle glucose uptake and whole-body glucose tolerance, all likely reflecting in utero programmed/unmasked impaired muscle function.

Skeletal muscle extracellular matrix (ECM) provides a framework structure that holds myofibers, blood capillaries and nerves supplying the muscle [68] and alterations in components of the ECM are associated with muscle dysfunction and impaired insulin actions [17]. Impaired capillary density in muscle is apparent in diabetes in both lean and obese rodents [69,70]. In addition, offspring of adverse suboptimal maternal nutrition have demonstrated endothelial dysfunction and remodeling of skeletal muscle vasculature, with limited microvascular and vasodilatory reserve [71]. Furthermore, there is evidence that IUGR affects collagen content in skeletal muscle [72]. Interestingly, in the current study in early adulthood, a significant diminished number of arterioles per myofiber area in LBW guinea pigs was observed, irrespective of the postnatal diet. Furthermore, skeletal muscle of the LBW offspring also displayed increased interstitial collagen deposition per myofiber area, although not significant, highlighting a potential role of UPI-induced IUGR/LBW in modulating skeletal muscle fibrosis in early adulthood. These findings suggest a deterioration of structural properties of UPI-induced IUGR/LBW skeletal muscle into young adulthood which could decrease vascular delivery compound such as insulin within the muscle skeletal muscle [11] with aging.

LBW in humans is associated with glucose intolerance and insulin resistance in skeletal muscle and whole body in the young [73–75] and excessive energy intake may be regarded as a promoter of insulin resistance in humans [76]. In the current study alterations in whole-body glucose tolerance or skeletal muscle glucose uptake were not observed, however, UPI-induced IUGR/LBW was associated with an increase in total PKCε protein in the gastrocnemius muscle. Overexpression of PKCε in skeletal muscle precedes the onset of hyperinsulinemia and hyperglycemia and data suggest this overexpression may be causally related to the development of insulin resistance, possibly by increasing the degradation of insulin receptors [77]. The mechanisms underlying the persistent postnatal change in PKCε at 4 months of age in UPI-induced IUGR/LBW are not yet clear. Nevertheless, such alterations in PKCε protein in conjunction with the significant diminished number of arterioles per myofiber area in muscle of LBW offspring could signal the beginning of insulin resistance pathogenesis. Indeed, skeletal muscle endothelial/microvascular dysfunction is linked to the insulin-resistant state [71,78], whereby an altered capillary network and reduced diffusing capacity and hemodynamic regulation of vessels in skeletal muscle is observed [79].

Protein kinase B/AKT plays a prominent role in mediating many of the metabolic effects of insulin [80]. The protein kinase PKCs, PKC-θ and PKCε provide a negative regulation for AKT phosphorylation at Ser^473^ and Thr^308^ [81–83], which impairs AKT activity. IRS-1 phosphorylation of Ser^302^ and Ser^307^ is increased in insulin-resistant mice; furthermore, JNK1-mediated disruption of insulin receptor/IRS-1 interaction is dependent on IRS-1 phosphorylation at Ser^302^ and Ser^307^ [84]. IRS-1 is also phosphorylated at Ser^307^ via several mechanisms, including insulin-stimulated kinases or stress-activated kinases like IKKβ, which inhibits insulin signal transduction and contribute to peripheral insulin resistance [85,86]. In the current study a reduction in the phosphorylation of AKT at Ser^473^ and Thr^308^ and increase in total protein levels of PKCθ, PKCε, phosphorylation of IRS-1 at Ser^302^ and IKKβ at Ser^176/180^ were associated with WD intake. An increased IRS1 Ser^302^ phosphorylation in LBW/WD compared to NBW/WD offspring was also observed, highlighting potentially an exacerbated insulin signaling defect in UPI-induced IUGR/LBW offspring consuming the WD. Consistent with our observations, defects in skeletal muscle insulin signaling including reduced phosphorylation in AKT at Ser^473^, Thr^308^, IRS-1 Ser^302/307^ phosphorylation, IKKβ Ser^177/181^ phosphorylation, and increased total protein level of PKCθ have also been reported in skeletal muscle of rats fed a high fat diet [87,88]. Interestingly and of note, in human and animal studies, young LBW offspring also display altered PKC and AKT signaling pathways and these changes occurred in conjunction with maintenance of whole-body glucose tolerance [89,90], similar to what we have observed in our LBW and WD cohorts. It has been postulated that changes in expression and phosphorylation of the insulin signaling intermediates precedes the development of overt IR and glucose intolerance [90], providing a molecular signature associated with early stages, or pre-clinical status, of disease progression [91]. Therefore, the predominantly diet associated changes in PKCs levels, AKT, IKKβ and IRS1 phosphorylation status, in conjunction with the maintained glucose muscle glucose uptake, and the increased PKCε and GLUT5 protein in LBW highlight that at this young age, both WD and UPI-induced IUGR/LBW offspring present an early-stage of insulin resistance pathogenesis resembling a pre-diabetic state as described in other reports [92].

Increasing dietary saturated fatty acid (SFA) consumption induces insulin resistance with concomitant increases in muscle diacylglycerol (DAG); and diets rich in n-6 polyunsaturated fatty acids appear to prevent insulin resistance by directing fat into triglycerides (TG), rather than other lipid metabolites [93]. In the current study, increased levels SFAs including myristic fatty acid were observed in skeletal muscle of WD-fed offspring and reductions in the omega-6 and omega-3 poly-unsaturated fatty acids in the neutral lipid fraction, which have also been implicated in reduced insulin sensitivity [93,94]. The alterations in fatty acids levels observed in the skeletal muscle of WD-fed offspring, appeared to mirror the composition of the dietary fats consumed in the WD [95], and may contribute to the alterations in AKT, IRS1, and IKKβ phosphorylation and PKC levels noted in these offspring. Indeed, increased availability of fatty acids has been linked to skeletal muscle insulin resistance and changes in key insulin signaling components [96,97].

The muscle fatty acid profile was however unaltered in the LBW CD offspring muscle, prompting further investigation into mitochondrial and amino acid metabolism readouts, as an alternative theory to explain the observed increased PKCε in muscle of LBW offspring because overexpression of PKCε is causally related to the development of insulin resistance, possibly by increasing the degradation of insulin receptors [77]. Interestingly, several medium- and long chain acylcarnitines were significantly increased in LBW CD or WD offspring independently. Acylcarnitine derangements which are similar to our results have been previously described in human IUGR cord blood samples at birth [28], in skeletal muscle of high-fat-fed mice [6] and in serum of prediabetic and diabetic patients [32]. Our results also showed an interactive effect of LBW and postnatal WD consumption on accumulation of medium and long chain acylcarnitines; a finding similar to data that was previously reported in plasma of protein restriction-induced LBW rats fed a high-fat/high sucrose-maltodextrin diet during adulthood for 10 weeks [98]. Altogether, these data shed light on unique muscle metabolic derangements in the young adult LBW CD and WD fed offspring..

Acylcarnitines arise from the conjugations of acyl-coenzyme A with carnitine for the transport of long-chain fatty acids across the inner mitochondrial membrane for *β*-oxidation and skeletal muscle is thought to be a principal contributor to the serum acylcarnitine pool [6]. Acylcarnitines are known to accumulate in situations of incomplete fatty acid oxidation and mitochondrial overload, where they are indicative of impairment of substrate flux through the TCA cycle and oxidative phosphorylation [6,99,100]. For example, chronic overnutrition leads to fatty acid oxidation rates that outpace the tricarboxylic acid cycle (TCA). This imbalanced environment exacerbates incomplete β-oxidation and leads to intramitochondrial accumulation of acyl-CoAs and respective acylcarnitines, a situation termed mitochondrial overload [6,99]. Carnitine Palmitoyltransferase CPT1b, is the mitochondrial enzyme that catalyzes the first and essential step in β-oxidation of long-chain fatty acids [101], and in the current study *CPT1b* mRNA was reduced in LBW offspring, irrespective of postnatal diet, indicating a potential defect in β-oxidation as indicated in the soleus of IUGR rats [102]. Additionally, 3-ketoacyl-CoA thiolase *(KT)* mRNA, the enzyme responsible for catalyzing the removal of acetyl-CoA from the fatty acid chain [101], was also reduced in LBW offspring. These indicators of defect in the oxidative process would slow down the rate of β-oxidation and impair substrate utilization through the remaining steps of the oxidative process, preventing complete oxidation of the fatty acid chains. However, the reduced levels in the expression of these fatty acid catabolism and oxidative genes were LBW-specific, highlighting that the accumulation of acylcarnitines in the WD-fed offspring are likely due to alternate defects in other components along the oxidative pathway. Indeed, palmitate in phospholipids was decreased in WD offspring and, the top components of our principal component analysis were composed primarily of medium and long-chain acylcarnitines: that explained the greatest sample variance more than any other factor of the model, with a significant main effect of WD observed in this cluster. As such, it is possible that the defects inducing acylcarnitine accumulation in LBW offspring versus WD-exposed offspring occur at different levels of the fatty acid catabolism process, namely β-oxidation or oxidative phosphorylation in LBW offspring, and TCA cycle in the WD-exposed offspring. These differential mechanisms, when superimposed upon one another in the LBW/WD offspring would have the potential to impair the oxidative process to a greater extent, promoting increased acylcarnitine accumulation, similar to the interactive effect observed in C8, C10, C12, C14:1, C16 and C18 acylcarnitine levels. As such, postnatal consumption of a WD following a UPI-induced IUGR/LBW may exacerbate mitochondrial overload, promoting increased accumulation of acylcarnitine intermediates.

Accumulation of acylcarnitines and amino acids may be at play early in diabetic progression, along with changes in DAG content and ceramide lipid intermediates [18,32,103]. Indeed, studies have highlighted that long-chain acylcarnitines interfere with muscle insulin signaling at the level of AKT phosphorylation and are associated with muscle insulin resistance [7]. Moreover, altered insulin sensitivity in the absence of DAG accumulation has been observed in pre-diabetic men [104]. Consistent with these previous reports, we did not observe any significant accumulation of triglycerides or DAG in the skeletal muscle of WD offspring. Taken together, the increase accumulation of short and long-chain acylcarnitines in LBW and WD offspring suggest such changes as predictive of impaired muscle insulin signaling markers, namely increased PKCs, increased IKKβ and IRS1 phosphorylation and reduced AKT phosphorylation in these offspring.

Amino acids are the basic units constituting proteins; they can maintain nitrogen balance, can be transformed into sugar or fat, and are involved in the structures of enzymes, hormones, and some vitamins [105]. In the present study, significant reductions in Glu and trend of increased Leu in skeletal muscle of LBW offspring irrespective of WD were identified. In the UPI-induced IUGR sheep model, fetuses display reduced net uptake rates of essential amino acids including Glu and branch chain amino acids (BCAAs: Leu, Val and Iso), Lys, Thr, Phe, His, and Trp in hindlimb muscle compared to control fetuses [26]. Additionally, research suggests that serum Glu levels in nascent MetS were significantly decreased compared to controls [18] and Glu can be used as an alternative energy source in patients with metabolic disorders [106]. Thus, decreased Glu in skeletal muscle of LBW guinea pig offspring suggest this BCAA is being used as an alternative energy source, leading to its depletion.

The amino acids, Ala, Asp and Phe were also significantly elevated in skeletal muscle of WD-fed offspring, an interesting observation, given high serum levels of these amino acids namely Ala and Phe, are found in prediabetes or MetS [106]. Ala is a glucogenic amino acid that can facilitate glucose metabolism, help alleviate hypoglycemia, and improve the body’s energy [105], similar to Asp which is also a glucogenic amino acid and necessary for protein synthesis in mammalian cells [107]. Additionally, Phe is a glucogenic/ketogenic amino acid used to produce proteins and important molecules, and elevated Phe levels are associated with severe metabolic disturbance [108]. Previous observations of the impact of postnatal exposure to high-fat diet report a reorganization of amino acid metabolism involving tissue-specific effects, and in particular a decrease in the relative allocation of amino acid to oxidation in several peripheral tissues including tibialis anterior muscle of young adult rats [109]. As such, one could speculate that the WD offspring in the current study, had a reduction in the relative allocation of certain amino acids to oxidation in skeletal muscle that may contribute to their elevation, highlighting potential amino acid oxidation defects, which should be further investigated. Finally, associations of changes in plasma and muscle BCAA (Val, Leu/Iso) and other amino acids (Phe, Tyr, Ala) with muscle insulin resistance, whole body insulin resistance and type 2 diabetes have been extensively reported and subsequently demonstrated with broad-based metabolomic investigations [8,18]. Taken together, the increase accumulation of certain amino acids in LBW and WD offspring may also be related to impaired muscle PKC levels, and IKKβ, IRS1 and Akt phosphorylation in these offspring given that BCAAs, particularly Leu, inhibit early steps in insulin action critical for glucose transport, including decreased insulin-stimulated tyrosine phosphorylation of IRS-1 and IRS-2, and a marked inhibition of insulin-stimulated phosphatidylinositol 3-kinase [10].

## 5. Conclusions

IUGR culminating in LBW and postnatal WD consumption are both independently and in conjunction, associated with diminished arteriole density, mitochondrial lipid metabolism and markers of insulin signaling in young adult lean offspring. Certainly, fetal hypoxia subsequent to UPI is associated with promoted oxygenated blood flow to the heart and reduced umbilical blood supply to organ such as skeletal muscle, an adaption which is understood to reprogram muscle mitochondrial lipid metabolism *in utero* likely through altered epigenetic regulation [110]. The current report further highlights that consumption of WD following UPI-induced IUGR/LBW likely unmasks LBW-induced evidence of metabolic dysfunction, exacerbating skeletal muscle acylcarnitine accumulation and promoting mitochondrial overload. These persistent alterations in acylcarnitine accumulation, and associated changes in the skeletal insulin signaling occur early in postnatal life, without body composition changes and weight gain and overt disruption of whole-body glucose tolerance, suggestive of a pre-diabetic situation in these male offspring.

## Author Contributions

K.D., O.S., J.A.T. and T.R.H.R. designed the experiments. K.D., O.S., N.S., L.Z., K.N., J.A.T., J.H., N.B., T-Y.L. and T.R.H.R. performed the experiments. K.D., O.S. N.S. and T.R.H.R analysed data and drafted the manuscript. Y.B. contributed to statistical analysis. All authors reviewed and edited the manuscript.

## Institutional Review Board Statement

The study was conducted according to the guidelines of the Declaration of Helsinki, and Animal care, maintenance, and surgeries were conducted in accordance with standards and policies of the Canadian Council on Animal Care, and the Western University Animal Care Committee reviewed, approved and monitored all procedures (Protocol #AUP-2009-229).

---

**Supplementary Table A1.**
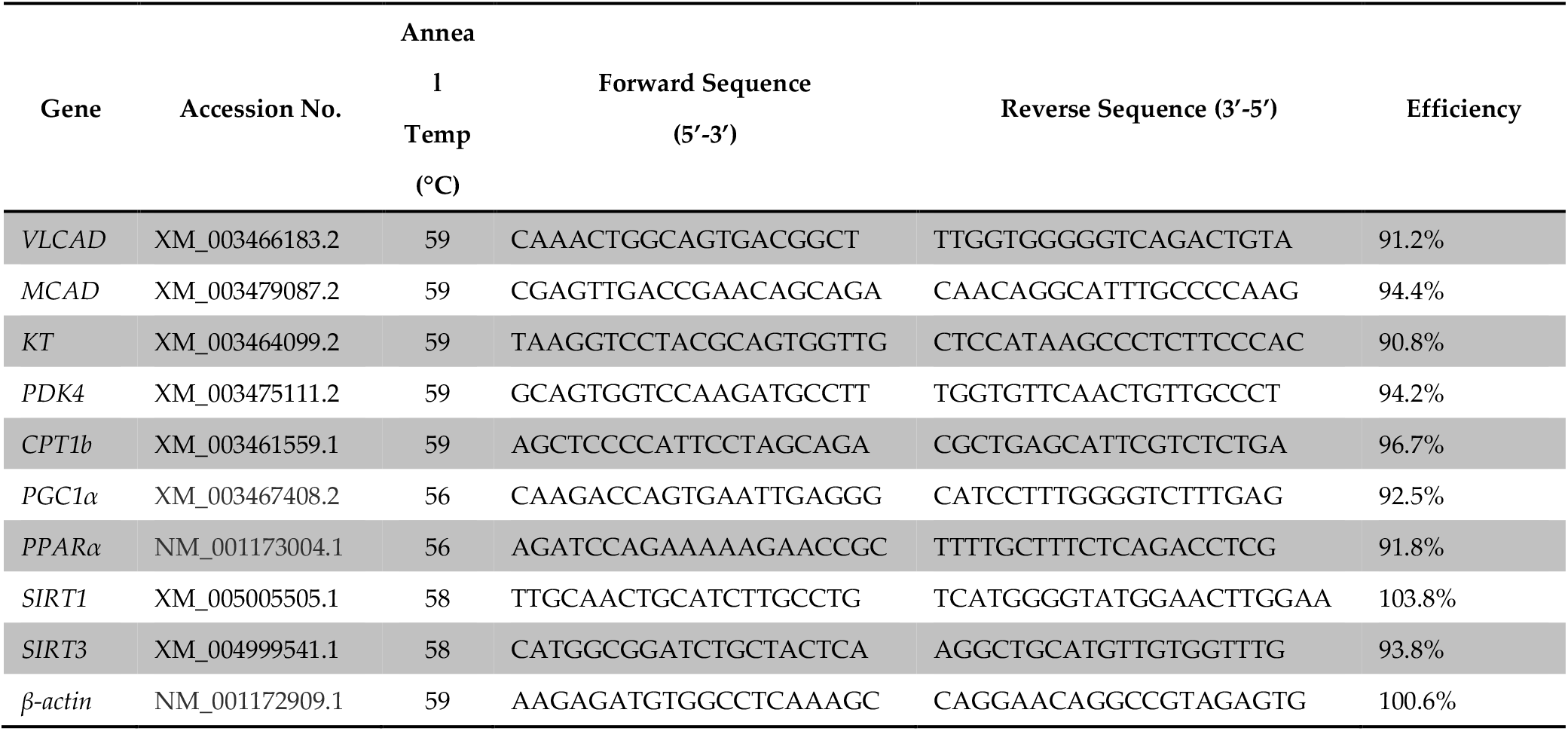
Primers used for analysis of gene expression by qRT-PCR.

**Supplementary Table A2.** Specifications and catalog numbers of antibodies used for immunoblotting.

| Protein | Species | Dilution | Blocking | Company | Catalogue No. |
| --- | --- | --- | --- | --- | --- |
|  |  |  | Solution |  |  |
| IRβ | Rabbit Monoclonal | 1:1000 | 5% BSA | Cell Signaling Technology | 3025 |
| pIRβ (Tyr 1150/1151) | Rabbit Monoclonal | 1:1000 | 5% BSA | Cell Signaling Technology | 3024 |
| pAKT (Ser 473) | Rabbit Monoclonal | 1:1000 | 5% BSA | Cell Signaling Technology | 4060 |
| pAKT (Thr308) | Rabbit Monoclonal | 1:1000 | 5% BSA | Cell Signaling Technology | 2965 |
| PKCθ | Rabbit Polyclonal | 1:1000 | 5% BSA | Cell Signaling Technology | 2059 |
| PKCε | Rabbit Monoclonal | 1:1000 | 5% BSA | Cell Signaling Technology | 2683 |
| pJNK(Thr183/Tyr185) | Mouse Monoclonal | 1:500 | 5% Milk | Santa Cruz Biotechnology | sc-6254 |
| JNK | Mouse Monoclonal | 1:500 | 5% Milk | Santa Cruz Biotechnology | sc-7345 |
| pIKKβ (Ser177/181) | Rabbit Polyclonal | 1:1000 | 5% BSA | Cell Signaling Technology | 2694 |
| IKKβ | Rabbit Polyclonal | 1:1000 | 5% BSA | Cell Signaling Technology | 2678 |
| pIRS1 (ser302) | Mouse Monoclonal | 1:1000 | 5% Milk | Millipore | 05-1086 |
| IRS1 | Rabbit Monoclonal | 1:1000 | 5% BSA | Cell Signaling Technology | 2382 |
| Anti-Rabbit Secondary | Mouse Monoclonal | 1:10,000 | 5% BSA or Milk | Cell Signaling Technology | 7074 |
| Anti-Mouse Secondary | Donkey | 1:5000 | 5% BSA or Milk | Cell Signaling Technology | 7076 |

